# Complex structural variation and behavioral interactions underpin a balanced sexual mimicry polymorphism

**DOI:** 10.1101/2024.05.13.594052

**Authors:** Tristram O. Dodge, Bernard Y. Kim, John J. Baczenas, Shreya M. Banerjee, Theresa R. Gunn, Alex E. Donny, Lyle A. Given, Andreas R. Rice, Sophia K. Haase Cox, M. Luke Weinstein, Ryan Cross, Benjamin M. Moran, Kate Haber, Nadia B. Haghani, Jose Angel Machin Kairuz, Hannah R. Gellert, Kang Du, Stepfanie M. Aguillon, M. Scarlett Tudor, Carla Gutierrez-Rodríguez, Oscar Rios-Cardenas, Molly R. Morris, Manfred Schartl, Daniel L. Powell, Molly Schumer

## Abstract

How phenotypic diversity originates and persists within populations are classic puzzles in evolutionary biology. While polymorphisms hypothesized to be under balancing selection segregate within many species, it remains rare for the genetic basis and the selective forces to both be known for the same trait, leading to an incomplete understanding of many classes of polymorphisms. Here, we uncover the genetic architecture of a balanced sexual mimicry polymorphism and identify behavioral mechanisms that may be involved in its maintenance in the swordtail fish *Xiphophorus birchmanni*. We find that ∼40% of *X. birchmanni* males develop a “false gravid spot”, a melanic pigmentation pattern that mimics the “pregnancy spot” associated with sexual maturity in female live-bearing fish. Using genome-wide association mapping, we detect a single intergenic region associated with variation in the false gravid spot, which is upstream of *kitlga*, a gene involved in melanophore patterning. By performing long-read sequencing within and across populations, we identify complex structural rearrangements between alternate alleles at this locus. The false gravid spot haplotype drives increased allele-specific expression of *kitlga*, which provides a mechanistic explanation for the increased melanophore abundance that causes the spot. By studying social interactions in the laboratory and in nature, we find that males with the false gravid spot experience less aggression; however, they also receive increased attention from other males and are disdained by females. These behavioral interactions may play a role in maintaining this phenotypic polymorphism in natural populations. We speculate that structural variants affecting gene regulation may be an underappreciated driver of balanced polymorphisms across diverse species.

## Introduction

Understanding how diverse phenotypes arise and persist are longstanding goals in evolutionary biology^1,2^. While directional selection can be invoked to explain the endless forms that have evolved in different species as they adapt to unique environments, this mechanism does not explain the persistence of phenotypic and genetic variation within interbreeding populations^3^. Indeed, the maintenance of phenotypic polymorphisms is especially puzzling when trait variation is predicted to directly affect fitness, under the expectation that adaptive variants should quickly fix. In the short-term, intermediate frequency polymorphisms may reflect neutral genetic drift, an allele on its way to fixation, or demographic processes such as migration^4^. However, since directional selection and drift are expected to lead to the loss of diversity, cases where phenotypic polymorphisms persist over long timescales or at stable frequencies are thought to be indicative of balancing selection^3^. Balancing selection encompasses several distinct mechanisms that can maintain variation, including heterozygote advantage, frequency dependent selection, spatially or temporally variable selection, and some forms of antagonistic selection^3^.

Understanding the genetic architecture and selective mechanisms underpinning balanced polymorphisms have remained important goals of modern evolutionary biology.

On the genetic level, theory predicts that discrete phenotypes segregating in a population should have a simple genetic basis, where phenotypic variation is explained by one or a few genetic loci^5–7^. Empirical work has uncovered many instances where single loci underpin phenotypic polymorphisms, although this may be in part driven by low power and discovery biases^8,9^. However, despite mapping to single regions of genomes, the genetic basis of these polymorphisms can be diverse, ranging from single nucleotide changes^10^ to megabase-scale structural variation^11–15^. Large inversions in particular appear to frequently underlie the evolution of phenotypic polymorphisms across the tree of life by limiting recombination between coadapted loci^16,17^, allowing phenotypically complex morphs to be inherited as simple Mendelian loci. However, while existing work on the genetic architecture of phenotypic polymorphisms has focused on large structural variants, there is increasing evidence that smaller structural changes can play an important role in both adaptation^18^ and disease^19^, raising the possibility that smaller structural variants may also play important roles in the genetic architecture of phenotypic polymorphisms. Such structural variation has been difficult to identify and study due to limitations of short-read sequencing, but recent advances in long-read technologies make it possible to precisely interrogate these variants and link them to variation in phenotype^20,21^.

Examples of balanced polymorphisms where both the causal genetic variants and selective mechanisms are known can be especially informative in understanding the origin and maintenance of phenotypic variation. Most well-studied examples with known genetic and fitness effects relate to immune function^4,22,23^ and mating system types^24,25^. In recent years, researchers have made progress linking genetic variation to phenotypic variation and selection in nature^26–28^ by leveraging natural systems where large-scale genomics projects are possible and mechanisms of selection can be studied in the laboratory and field. Despite this, many classes of polymorphisms remain poorly understood. For instance, sexual mimicry polymorphisms, where males mimic females (or vice versa), are a charismatic class of phenotypic polymorphisms that have repeatedly evolved across the tree of life^29,30^. Decades of behavioral and physiological studies have led to the expectation that sexual mimicry polymorphisms may be maintained by balancing selection, but in most cases they remain poorly characterized at the genetic level^30^. Sexual mimicry can manifest in various morphological^31^, chemical^32,33^, or behavioral^34^ traits, with selective mechanisms associated with alternative reproductive tactics^29^, reduced harassment^31^, and even physiological adaptation^35^. Due to the diversity of phenotypes and selective mechanisms acting on these traits, sexual mimicry polymorphisms present exciting opportunities to characterize the genetic, evolutionary, and population-level processes that contribute to the maintenance of genetic and phenotypic variation in natural populations.

Swordtail fish in the genus *Xiphophorus* are classic models of sexual selection driving the evolution of ornamentation and an excellent system for studying the genetic architecture and selective forces underpinning polymorphic traits^36–39^. Male *Xiphophorus* have evolved elaborated fin morphology and pronounced coloration, which are generally associated with increased reproductive success^38,40,41^. Additionally, female *Xiphophorus* develop a gravid spot at sexual maturity, which becomes more pronounced when the female is carrying mature eggs and embryos (since these develop internally)^42^. Enlarged gravid spots have been shown to increase male courtship incidence in some *Xiphophorus* species and are hypothesized to be a signal that males use to direct courtship^43^. In several *Xiphophorus* species, some males develop a superficially similar melanic pigmentation trait, called the “false gravid spot” (Fig. 1a). This phenotypic polymorphism was first described nearly a century ago^44^ and has long been presumed to be involved in sexual mimicry^45^. While the presence of a polymorphism shared by multiple related species is often thought to be a sign of balancing selection^46,47^, little is known about the selective forces acting on the false gravid spot, with only one prior study having investigated behavioral factors that may favor the trait^48^. Moreover, the genetic basis and developmental progression of this sexual mimicry polymorphism has not been described in any species.

**Figure 1.**
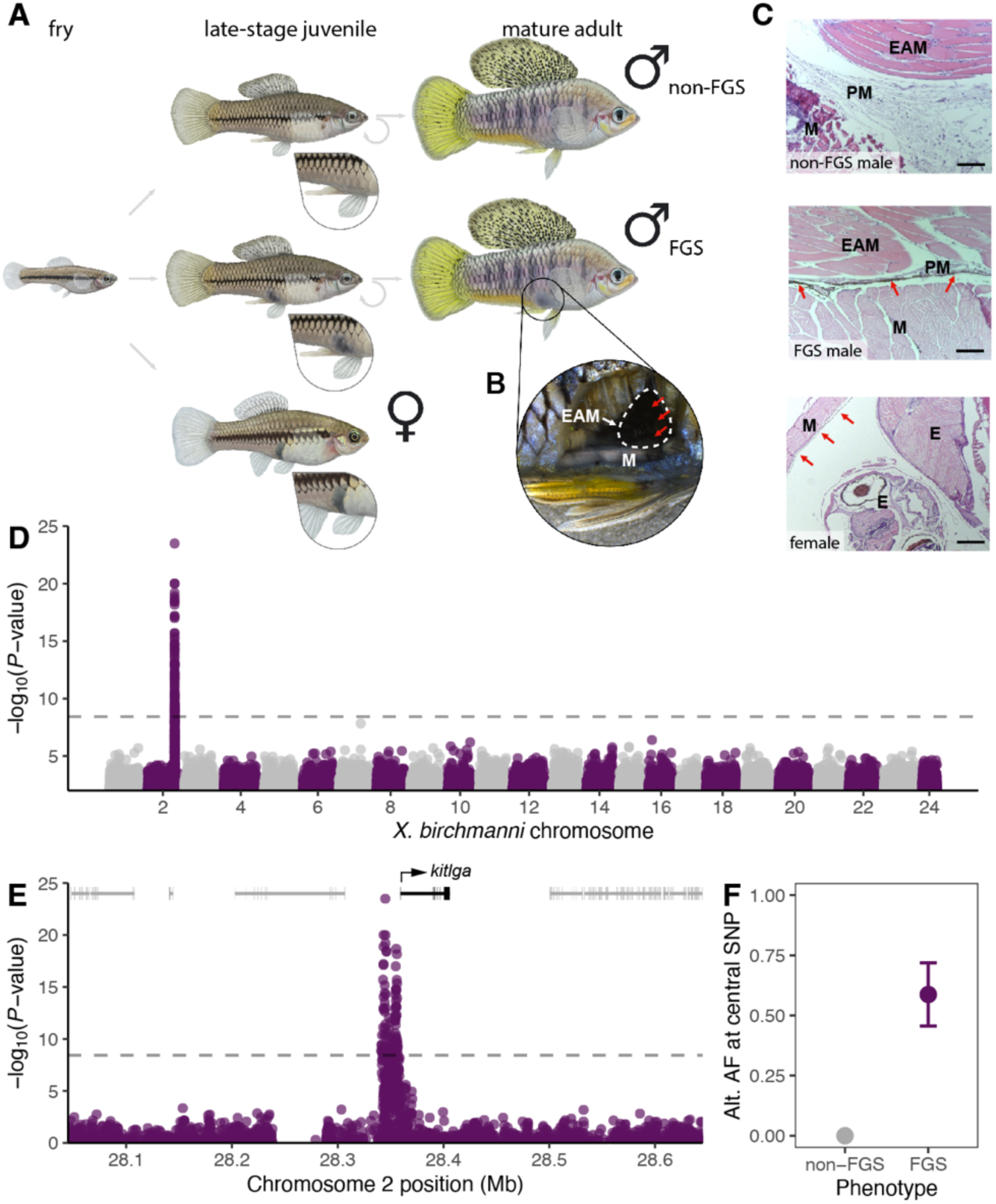
**The false gravid spot is a pigmentation polymorphism in *X. birchmanni* with a simple genetic basis. A**) The false gravid spot (FGS) is a sexual mimicry trait present in male *X. birchmanni*. Fry (left) are morphologically indistinguishable by sex until the onset of puberty. Upon sexual maturity, females (bottom) develop a gravid spot, which becomes larger and darker when carrying mature eggs or developing embryos. Non-false gravid spot (non-FGS) males develop no pigmentation in this region (top), while false gravid spot (FGS) males develop pigmentation anterior to the gonopodium (middle). The false gravid spot remains visible throughout the adult male’s life, even after sexual maturity and the development of other secondary sexual traits. Illustrations by Dorian Noel. **B**) Dissection of a false gravid spot male reveals the spot stems from pigmentation in the tissue surrounding the gonopodial suspensorium, for instance in the erector analis major (EAM) muscle, which is visible through the body wall musculature (M). **C**) Top: Histological sections indicate *X. birchmanni* males without the false gravid spot do not develop pigmentation in the erector analis major or its perimysium. Middle: *X. birchmanni* males with the false gravid spot accumulate melanophores in the perimysium of the erector analis major. Bottom: Histological sections confirm the gravid spot in *X. birchmanni* females is due to expansion of the highly pigmented peritoneum. Labels: M - body wall musculature; EAM - Erector analis major; PM - Perimysium of EAM; E - Embryo; red arrows - pigmented melanophore cells. Scale bars denote 50 µm for top and middle images and 200 µm for bottom image. **D**) Manhattan plot showing results from a genome-wide association study using 329 *X. birchmanni* males with 39% false gravid spot frequency. A single autosomal region is associated with the false gravid spot on the distal end of chromosome 2. Dashed line shows the 5% false positive rate determined by simulations. **E**) Manhattan plot highlighting the genome-wide significant region on the end of chromosome 2. The significant region spans 17 kb and does not overlap with any genes but is 1.5 kb upstream of the known pigmentation gene *kitlga* (black gene model). **F**) Allele frequency (AF) of the non-reference allele at the central representative SNP from the GWAS region (position 28349032) in non-FGS and FGS males. Error bars denote ± 2 binomial standard errors.

Here, we find that variation in the false gravid spot in *X. birchmanni* is controlled by a non-coding region upstream of *kitlga*, a gene that underlies pigment pattern variation in many vertebrates. The haplotypes associated with the false gravid spot display considerable structural complexity and drive higher allele-specific expression of *kitlga*. We demonstrate that the false gravid spot emerges in a male-specific tissue and develops before other male secondary sexual traits. Several lines of phenotypic and genetic evidence suggest that this polymorphism is maintained by balancing selection in nature. In laboratory behavioral trials and observations in the wild, we find the false gravid spot influences social interactions including aggression and courtship, hinting that behavioral interactions may contribute to the maintenance of this sexual mimicry polymorphism.

## Results

### The false gravid spot occurs in a male-specific tissue and is distinct from the female gravid spot

The gravid spot in female *Xiphophorus* arises during sexual maturation, when the heavily melanized peritoneal lining surrounding the internal organs expands during oogenesis and becomes visible through the body wall, forming an externally visible melanic spot^42^. While the false gravid spot in males also arises during puberty and localizes below the dermis^44^, it appears above the male gonopodium (the modified anal fin used for internal fertilization in *Xiphophorus* and other poeciliid species), slightly posterior to the position of the gravid spot in females (Fig. 1a). This led us to hypothesize that the anatomical basis of the false gravid spot and true gravid spot may be distinct. Dissections in *X. birchmanni* revealed that pigmentation originated from the tissue surrounding the gonopodial suspensorium bone structure in false gravid spot males (Fig. 1b; Fig. S1). This tissue complex is posterior to the peritoneal lining and separated from all internal organs. Histological sections revealed melanophore accumulation in a thin perimysium surrounding the erector analis major muscle of males with the false gravid spot, while this tissue was unpigmented in males lacking the trait (Fig. 1c). The musculature in both the body wall and erector analis major was unpigmented in males with both phenotypes, suggesting that the melanized perimysium was responsible for the external phenotype in *X. birchmanni*. We confirmed that the peritoneal lining is also melanized in females, (Fig. 1c), but females do not have an equivalent tissue structure to the perimysium of the erector analis major, demonstrating that the anatomical basis of the spot differs between males and females.

### Variation in the false gravid spot phenotype maps upstream of kitlga

To characterize the genetic basis of the false gravid spot polymorphism in *X. birchmanni*, we performed a case-control genome-wide association study (GWAS), searching for allele-frequency differences between phenotype classes. Our sample consisted of 329 males from a single population (Río Coacuilco), with 39% false gravid spot frequency, which had previously been sequenced at low coverage genome-wide^36^. To map the short-read data, we assembled a highly contiguous and complete chromosome-level *X. birchmanni* reference genome from an individual without the false gravid spot (Supplementary Information). Our GWAS recovered a single peak on chromosome 2 (Fig. 1d) which surpassed the genome-wide significance threshold estimated using simulations (p<3.7 × 10^-9^). This association remained strongly significant after correcting for population structure (always p<1 × 10^-10^; Fig. S2, Fig. S3). Chromosome 2 is expected to be an autosome in *X. birchmanni*, based on sex chromosome architecture in related species^36,49^.

The region associated with the false gravid spot spanned 17 kb and did not include any genes, but the closest annotated gene was *kit-ligand a* (*kitlga*, Fig. 1e), a gene with a well-documented and conserved role in melanocyte development and homeostasis across vertebrates^50,51^. We found the proximal end of the associated region fell 1.5 kb upstream of the likely *kitlga* transcriptional start site (Fig. 1e, Fig. S4). Within the significant region, we detected strong allele frequency differences between phenotype groups (Fig. 1f), consistent with a dominant and highly penetrant architecture of the false gravid spot. Analysis of the amino acid sequences of *kitlga* across multiple individuals of *X. birchmanni* sequenced at high coverage revealed no variation in amino acid sequence (Fig. S5), confirming that variation in the trait is not driven by coding differences. Moreover, comparisons to *X. birchmanni’*s sister species, *X. malinche,* indicated that rates of protein evolution in *kitlga* were consistent with strong purifying selection (dN/dS = 0.001; Fig. S6).

### A complex structural rearrangement is associated with the false gravid spot

To perform an initial investigation into haplotype structure in the associated region, we calculated linkage disequilibrium (LD) using 23 individuals previously sequenced at higher depth from the same population (median 18.4×). We found strong linkage disequilibrium (LD) between SNPs in the GWAS peak (Fig. 2a), with a rapid decay on either side of the region. To explore if this LD pattern was driven by structural differences between the false gravid spot and non-false gravid spot haplotypes, we generated a PacBio HiFi assembly for the false gravid spot haplotype (Supplementary Information), and generated a pairwise alignment between the false gravid spot and non-false gravid spot haplotypes using MUMmer4^52^. We found evidence of complex rearrangements in the haplotype associated with the false gravid spot (Fig. 2a). This complex rearrangement precisely localized with the strongest signal in the GWAS upstream of *kitlga* and the region of high LD. To investigate possible effects of our choice of reference sequence on the GWAS signal, we repeated the analysis replacing chromosome 2 with a HiFi contig derived from a false gravid spot *X. birchmanni* haplotype and found our results to be qualitatively unchanged (Fig. S7).

**Figure 2.**
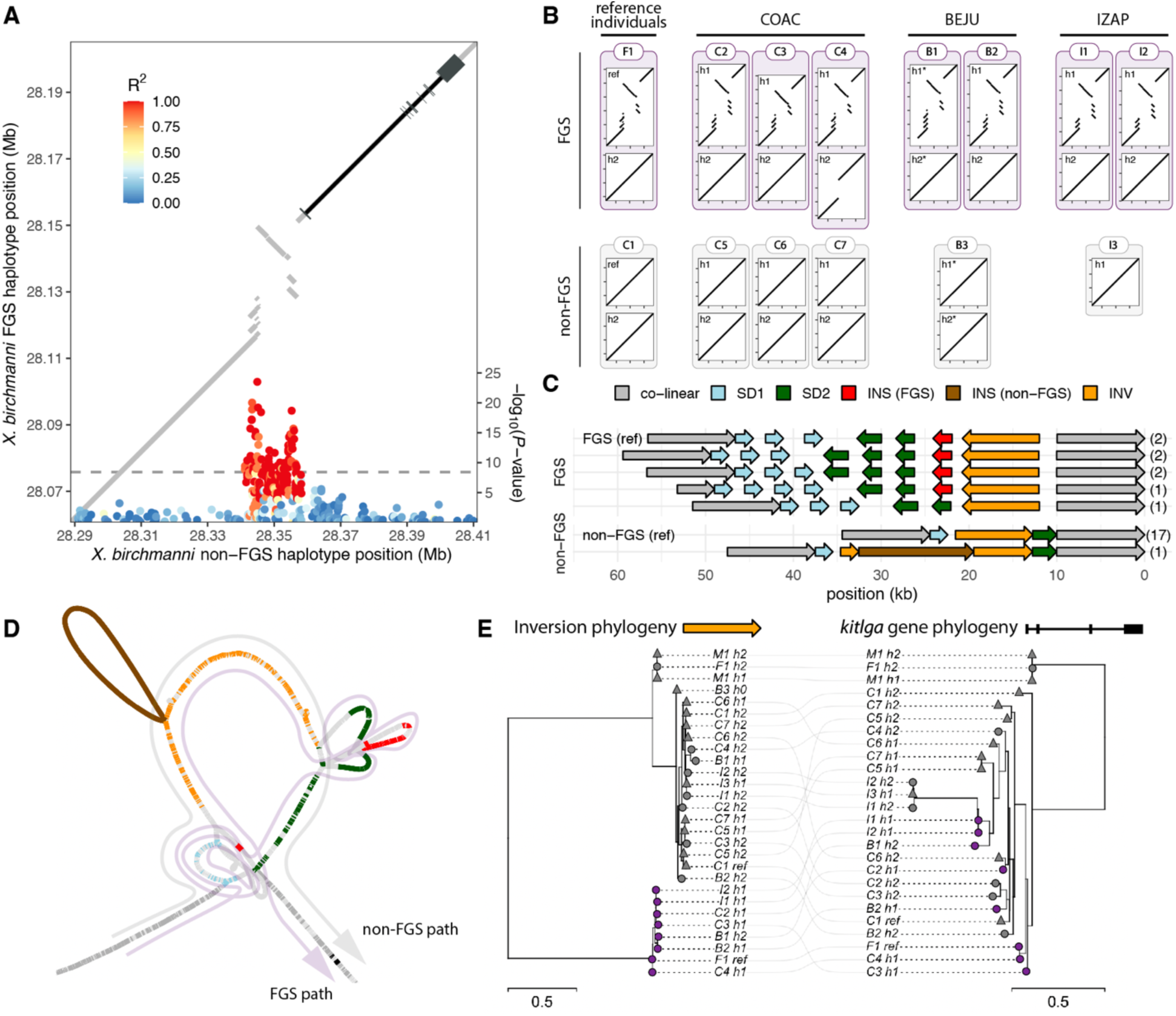
**A complex structural variant is associated with the false gravid spot locus. A**) Combined plot showing GWAS peak with SNPs colored by linkage disequilibrium (LD) to a central SNP overlayed with MUMmer4 alignment (gray lines) of false gravid and non-false gravid haplotypes. Color of individual points denotes R^2^ with the center SNP at position 28349032 bp. Region of high LD perfectly colocalizes with the complex structural variant in the MUMmer4 alignment, which contains two sets of segmental duplications, an insertion, and an inversion relative to the non-false gravid spot reference. The *kitlga* gene region is downstream of the structural variant. The gene region is superimposed on the alignment in black, with exons noted with thicker segments. **B**) Long-read sequencing of individuals from several *X. birchmanni* populations reveals that this complex structural variant is perfectly associated with the false gravid spot phenotype. Alignments between each haplotype compared to the reference sequence. Individuals with the false gravid spot phenotype are indicated in purple outlines and individuals without are noted in gray. Reference individuals include the F_1_ *X. birchmanni* × *X. malinche* hybrid with false gravid spot and the chromosome-level assembly from Coacuilco without false gravid spot. An additional 12 individuals were sequenced across 3 populations (COAC – Coacuilco, BEJU – Benito Juarez, IZAP – Izapa). Haplotypes were extracted from diploid assemblies, unless noted with a star, in which case individual long-reads spanning the region were used to infer haplotype structure. Note that the non-false gravid spot individual from Izapa (I3) was completely homozygous in this region, so a single haplotype was generated. **C**) Five structurally variable haplotype classes exist across 8 false gravid spot haplotypes sequenced, compared to a single structural variant across 17 non-false gravid haplotypes in *X. birchmanni*. Numbers on right denote number of times haplotype was observed. SD1 - segmental duplication 1; SD2 - segmental duplication 2; INS (FGS) - insertion in false gravid spot haplotype (piggyback 4 element); INS (non-FGS) - insertion in non-false gravid haplotype. **D**) Local variation graph reveals complexity in the regions surrounding the segmental duplications. Regions are colored as in **B**. The false gravid spot and non-false gravid spot haplotypes are represented by different paths through the graph. **E**) Local phylogeny of the inversion (left) versus *kitlga* gene sequence (exons and introns; right) in *X. birchmanni* and *X. malinche*, with light gray lines connecting the same haplotype. The inverted region clusters perfectly by false gravid spot phenotype, while the genic sequence clusters by species and not by phenotype. Haplotype labels same as in **B**. Circles denote whether the individual from which the haplotype was derived displayed a false gravid spot phenotype while triangles denote haplotypes from individuals with a non-false gravid spot phenotype. Colors denote if haplotype was inverted compared to the reference (purple) or syntenic to the reference (gray). Note that the false gravid spot haplotype is dominant (see main text), so individuals marked by circles can harbor both purple and gray haplotypes.

We next aimed to formally test if the structural variant was associated with the false gravid spot in *X. birchmanni*. Because of the length and complexity of the rearranged haplotype and shortcomings of short-read methods for identifying structural variation^53^, we generated *de novo* diploid assemblies for 12 additional individuals from 3 populations across the *X. birchmanni* species range using Oxford Nanopore Technologies (ONT) or PacBio HiFi long-read sequencing. For each haplotype, we generated a pairwise alignment compared to the *X. birchmanni* non-false gravid spot reference genome using MUMmer4 (Fig. 2b). In all *X. birchmanni* individuals with the false gravid spot phenotype (n=7), we identified one rearranged haplotype and one haplotype that was largely co-linear with the non-false gravid spot reference (Fig. 2b). In *X. birchmanni* individuals without the false gravid spot (n=7) all haplotypes were co-linear with respect to the non-false gravid spot reference (Fig. 2b). This result indicates that the false gravid spot haplotype is dominant with respect to phenotype. Moreover, all haplotypes from *X. malinche*, the sister species of *X. birchmanni* that lacks the false gravid spot, were also co-linear with the non-false gravid haplotype in *X. birchmanni* (Fig. 2b; Fig. S8). These results strongly connect the structurally complex rearrangement in *X. birchmanni* with the false gravid spot phenotype (p=0.001263, Fisher’s Exact Test). We note that while all wild-caught *X. birchmanni* with false gravid spot in our long-read dataset were heterozygous for the rearrangement, this is not unexpected given the sample size (p=0.3976 by simulation), and we detected 2 individuals homozygous for the false gravid spot allele in our larger high coverage short-read dataset.

Based on the pairwise alignments between reference individuals, the complex structural variant contained two distinct segmental duplications, an insertion, and an inversion (Fig. 2a; Fig. S9). Within the rearrangement, the two distinct duplications and the inversion displayed local identity (93-97%) between the false gravid and non-false gravid haplotypes (Fig. S9; outside of the rearrangement identity was >99.5%). In contrast, 2.5 kb of the insertion showed high similarity to a *Xiphophorus* piggybac protein 4 transposable element. Within the associated region, the false gravid spot haplotype also had a higher frequency and length of inverted repeats relative to the non-false gravid spot haplotype (Fig. S10).

Within haplotype classes, we noticed that some alignments displayed slight differences compared to the reference haplotypes (Fig. 2b). To compare haplotype architecture more quantitatively, we extracted the regions identified from the reference haplotypes (Fig. 2a) and used MUMmer4 to obtain regions of homology between all assemblies. We identified 5 distinct allele classes across 8 false gravid spot haplotypes sequenced (Fig. 2c; Fig. S11), which exhibited copy number variation in the distal segmental duplication (SD1; 3 vs 4 copies) and proximal segmental duplication (SD2; 2 vs 3 copies), presence of the insertion (0 vs 1 copy), and length of the region between the 2 duplications (∼1 kb vs 4 kb). In contrast to the high diversity of false gravid spot haplotypes, we only detected one structural variant haplotype (a distinct insertion) among the 18 non-false gravid spot haplotypes sequenced.

To better understand differences between haplotypes and overcome limitations of pairwise alignments in complex regions, we constructed a base-level variation graph (Fig. 2d) for this region using the pan-genome graph builder (PGGB)^54^. As expected, most elements identified in the pairwise analysis (e.g., the inversion) were represented once, with high copy number elements (e.g., SD1) in the graph traversed multiple times. The graph revealed additional topologies in regions that were not detected with pairwise alignment approaches (Fig. 2c), confirming the high level of complexity associated with the variable boundaries of haplotype variation in this locus.

We next aimed to explore the evolutionary history of the structural variant and the *kitlga* gene by constructing and comparing local phylogenies for each region. Based on the complexity uncovered by the alignments and variation graph in the structural variant, we focused on the 8.8 kb inversion, which was present as a single copy across all haplotypes. For each assembly, we extracted the inversion and the coding region of *kitlga* (including introns), and constructed local maximum likelihood phylogenies for each region using RaxML^55^ (Fig. 2e). In the alignment of the inverted region, we found the false gravid spot haplotypes and non-false gravid spot haplotypes formed distinct, monophyletic groups (Fig. 2e). Within the non-false gravid spot haplotypes, *X. birchmanni* and *X. malinche* haplotypes grouped separately. Taken together, this suggests the false gravid spot and non-false gravid spot haplotypes diverged prior to species divergence. However, the same was not true for the *kitlga* transcript sequence, which clustered by species but not by phenotype. Together, this phylogenetic analysis highlights a history of recombination between the structural variant and the coding sequence of *kitlga* and underscores the importance of the structural variant in producing the false gravid phenotype.

### Expression of kitlga in false gravid spot males

Given the localization of the structural rearrangement to a non-coding region and the lack of *kitlga* amino acid variation, we investigated if expression differences in *kitlga* might drive the false gravid spot phenotype. We performed mRNA sequencing in 13 juvenile males with and without false gravid spot, using pseudoalignment with kallisto^56^ and differential expression analysis with DESeq2^57^ (Table S1). *kitlga* was among the most differentially expressed genes genome-wide between false gravid spot and non-false gravid spot males in the erector analis major and its perimysium (Fig. 3a) but was not differentially expressed in brain (Fig. 3b). *kitlga* was also differentially expressed in the muscle tissue above the erector analis major and its perimysium, although the expression difference was smaller (Fig. S12). We also confirmed differential expression of *kitlga* in the erector analis major and its perimysium using quantitative real-time PCR (Fig. S13; Table S2). In RNA-seq analyses of the erector analis major and its perimysium, we also found increased expression of several other genes with known roles in melanogenesis pathways (Table S1), consistent with increased melanophore abundance in this tissue. However, *kitlga* was the only gene within 300 kb of the GWAS peak that was differentially expressed (Fig. 3c), suggesting it might act upstream of other differentially expressed genes identified genome-wide.

**Figure 3.**
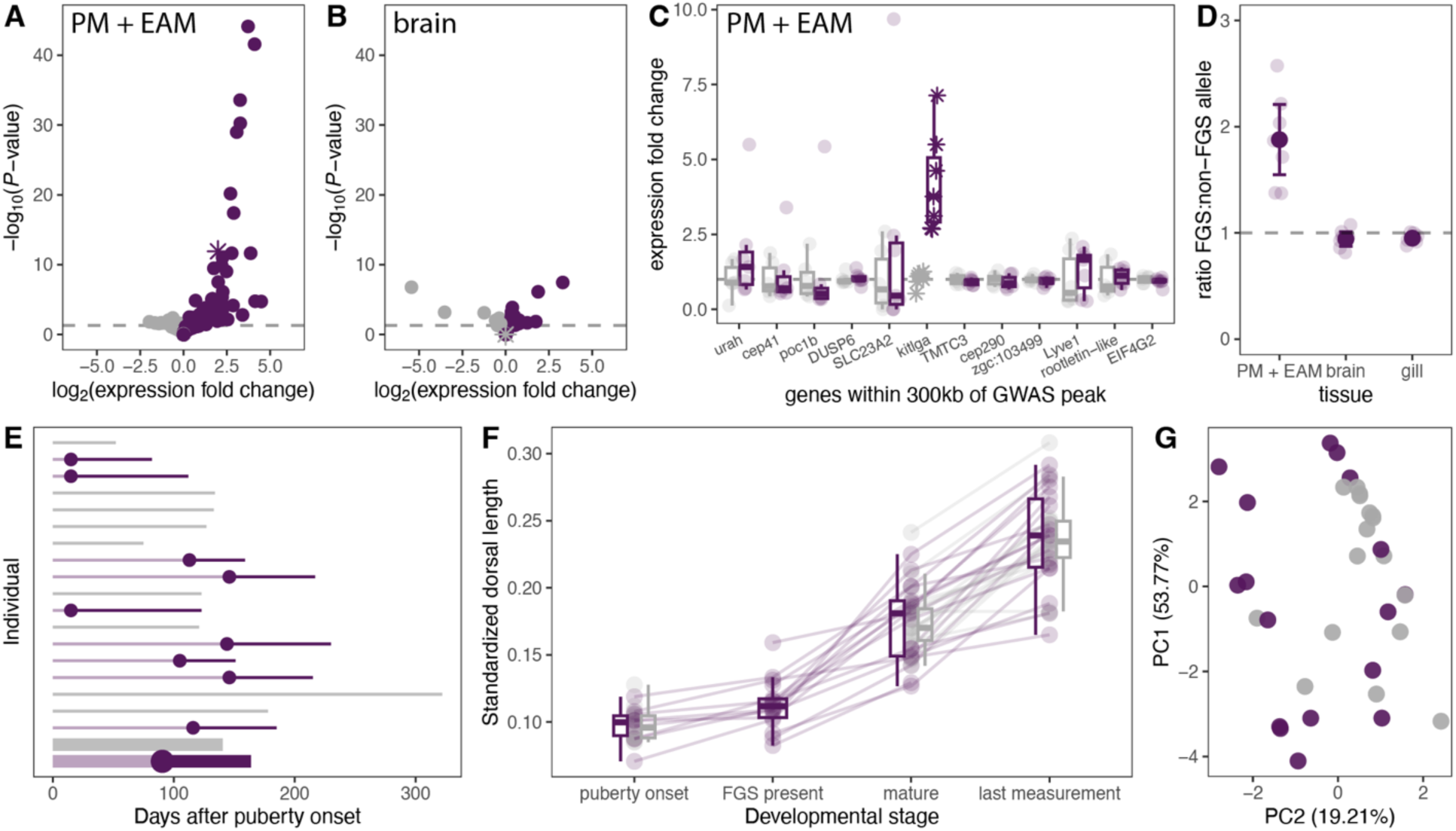
**Gene expression and developmental timing of the false gravid spot phenotype. A**) RNA-seq analysis shows differential expression of several genes in individuals with and without false gravid spot in the erector analis major tissue and its perimysium (EAM + PM). *kitlga* is shown with a star. Genes with increased expression in false gravid spot individuals are shown in purple and genes with increased expression in non-false gravid individuals are shown in gray. **B)** Brain tissue shows limited differential gene expression between false gravid spot and non-false gravid spot individuals, including for *kitlga*. **C**) In the combined EAM and PM tissue, *kitlga* is the only differentially expressed gene within 300 kb of the GWAS peak. Expression fold change of the 12 closest genes to the peak (ordered by genomic position), with non-false gravid individuals in gray and false gravid individuals in purple. Expression for each individual is normalized to mean expression of non-false gravid individuals, with boxplots showing data quartiles and points representing individuals. **D**) Allele-specific expression in *X. birchmanni* × *X. malinche* hybrids carrying a false gravid and non-false gravid allele, quantified by pyrosequencing, suggests the expression differences in *kitlga* are under *cis-*regulatory control. Expression is normalized to the *X. malinche* allele. Large points and whiskers denote mean ± 2 binomial standard errors, and small points represent individuals. **E)** Developmental time between onset of puberty until sexual maturity in 17 *X. birchmanni* males in the laboratory. Individuals are ordered from top to bottom by birth order, false gravid spot individuals are noted in purple, with the dot showing when the spot developed, non-false gravid individuals are in gray. Thick bars and dot and at the bottom represent mean time to sexual maturity and false gravid spot timing for each group. **F)** Dorsal fin length, standardized by body length, across fish from the developmental series shows that dorsal fin elongation tends to occur later during sexual maturity and continues to elongate after males are reproductively mature. Males with (purple) and without false gravid spot (gray) do not differ significantly in their dorsal length in laboratory conditions. **G**) PCA of male phenotypes at sexual maturity shows individuals with and without false gravid spot raised in laboratory conditions do not systematically differ in their secondary sexual traits. Traits scored include body and fin morphometrics, as well as pigmentation traits.

The position of the structural variant adjacent to the *kitlga* promotor suggests that it could modulate expression as a *cis*-regulatory element, a hypothesis we tested by measuring allele-specific expression. Given that few SNPs segregate in the *X. birchmanni kitlga* coding sequence, we measured allele-specific expression in laboratory hybrids with the false gravid spot, generated between *X. birchmanni* and *X. malinche*. These individuals necessarily inherit their false gravid spot haplotype from *X. birchmanni*. While the non-false gravid spot haplotype structure in *X. malinche* and *X. birchmanni* are similar (Fig. 2b, Fig. S8, Fig. S11), the *kitlga* coding sequence has accumulated several synonymous substitutions between species (Fig. S14), which allow us to distinguish whether a transcript originated from the false gravid spot (*X. birchmanni*) or non-false gravid spot (*X. malinche*) haplotype. Using a pyrosequencing approach (Table S3), we found strong differential expression of the false gravid and non-false gravid spot alleles in erector analis major and perimysium (Fig. 3d, Fig. S15; p<0.001) and little evidence of differential expression in other tissues (Fig. 3d). Together, this suggests that the structural variant drives increased allele-specific expression of *kitlga* in a tissue-specific manner.

### The false gravid spot develops before sexual maturity and before other secondary sexual traits

The developmental timing of sexual mimicry traits can help to inform hypotheses about their function^30^. To determine when the false gravid spot developed, we tracked the phenotypic progression of 43 males raised together in standard conditions from the onset of puberty through >1 year of age. We defined the onset of puberty as when males began to develop a gonopodium, and defined completion of sexual maturity as when the gonopodium became a functional intromittent mating organ^58–60^ (Fig. S16; Table S4). We found that males begin to develop pigmentation in the false gravid spot area coincidently with the early stages of external gonopodium development, in many cases several months before they complete sexual maturity (Fig. 3e). Thus, the results of this developmental series suggest that development of the false gravid spot, driven by pigmentation in the perimysium of the erector analis major (Fig. 1c), is temporally linked to external development of the gonopodium. This is expected anatomically given that modification of anal fin rays which form the gonopodium coincides with the development of the gonopodial suspensorium to which the erector analis major is anchored^61^.

Importantly, we also found that two sexually dimorphic traits, the dorsal fin and vertical bars, which are used for intra- and inter-sexual communication in *Xiphophorus*^62,63^, lagged the development of the false gravid spot. Most elongation of the dorsal fin occurred after false gravid spot development and continued after sexual maturity (Fig. 3f). However, lab-reared males with and without the false gravid spot did not differ significantly in overall ornamentation in adulthood (Fig. 3g). This suggests that, while the false gravid spot develops before sexual maturity, it is not developmentally linked to variation in other sexually dimorphic traits in adult fish raised under laboratory conditions.

### The false gravid spot shows evidence of being a balanced polymorphism

With a better understanding of the genetic and developmental basis of the false gravid spot, we sought to investigate if selection is acting on the trait in nature. We first measured the frequency of the false gravid spot phenotype across the *X. birchmanni* range, with the expectation that under some models of balancing selection, it would exhibit similar phenotypic frequencies across populations and not be fixed in any population. We sampled 9 *X. birchmanni* populations (542 total individuals) and found the false gravid spot was polymorphic in all locations, with observed phenotypic frequencies ranging from 0.11–0.50 (Fig. 4a). Furthermore, in populations with high confidence frequency estimates (≥100 individuals sampled), phenotypic frequencies ranged from 0.24-0.47. At Coacuilco, we repeatedly sampled frequency from 2017 to 2023, finding the phenotypic frequency fluctuated between 0.30 and 0.60 during this interval (Fig. S17), with some significant changes between years (two proportion Z-test, p=0.015).

**Figure 4.**
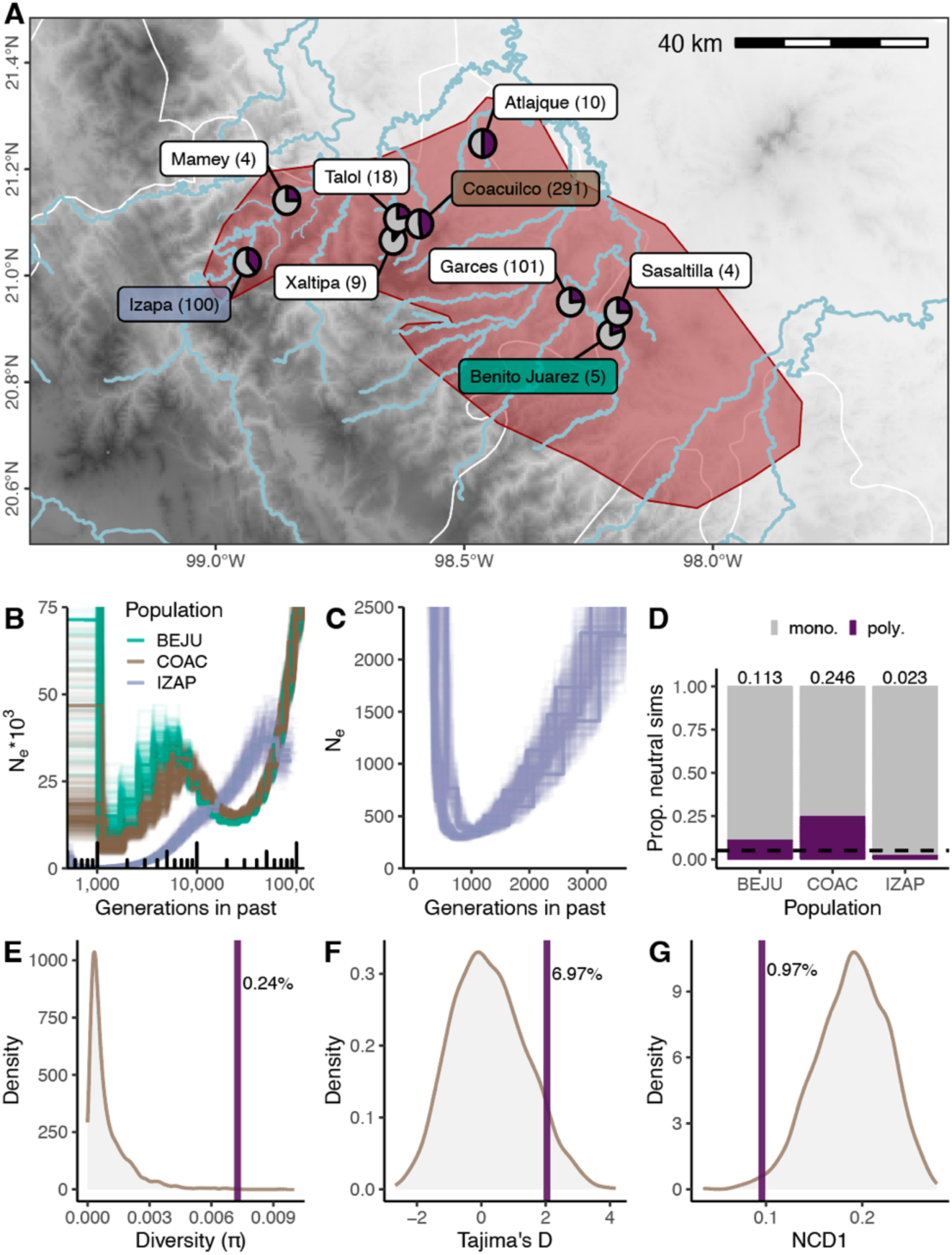
The false gravid spot is under balancing selection in the wild. A) The false gravid spot is present at all sampled populations across the *X. birchmanni* range (red polygon). Pie charts depict false gravid spot phenotypic frequency in adult males, ranging from 0.11 to 0.50, with sample size in parentheses. Map shows elevation from 0 (white) to 3000 meters (black), with major rivers shown in blue, and state boundaries shown in white. **B)** PSMC plot showing inferred population histories over the last ∼100k generations of three *X. birchmanni* populations, Benito Juarez (BEJU), Coacuilco (COAC) and Izapa (IZAP) with variants called from ONT data. Izapa experienced a distinct demographic history from Benito Juarez and Coacuilco, including a sustained bottleneck. **C)** Recent population history at Izapa inferred from PSMC, showing a minimum N_e_ of 290 individuals over ∼1000 generations. **D)** Outcome of simulations of neutral polymorphisms (10,000 simulations) under the demographic history inferred for each population from PSMC. In the absence of selection, polymorphisms are maintained in only 2.3% of simulations matching the demographic history in Izapa. **E)** Nucleotide diversity (pi) within the inverted region of the false gravid spot haplotype in 23 individuals at Coacuilco (purple line) compared to the distribution of all 8.8 kb windows from chromosome 2 (gray distribution). The pi value calculated in the inversion is in the top 0.24% chromosome-wide. **F)** Tajima’s D within the inversion (purple line) falls within the top 6.97% of values chromosome-wide. **G)**. The NCD1 value within the inversion (purple line) is in the bottom 0.97% of values chromosome-wide.

Since some of the *X. birchmanni* populations surveyed occur in different river drainages, we sought to better understand the demographic history of these populations. We leveraged our long-read data from three of these populations (Coacuilco, Izapa, and Benito Juarez), called variants, used PSMC^64^ to infer demographic history of each population (Fig. 4b), and estimated divergence between populations (Fig. S18). We discovered that the Izapa population underwent a sustained bottleneck (estimated N_e_ as low as 293 individuals) for several thousand generations, but still has a false gravid spot frequency of 39% (Fig. 4c). To explore how likely the maintenance of the false gravid spot would be in the absence of selection, we used SLiM^65^ to simulate a neutral locus in populations matching the observed demographic histories. In the simulated Izapa population, we found a neutral locus was only maintained as polymorphic in 2.3% of all neutral simulations (Fig. 4d; n=10,000 simulations). This indicates that, under neutrality, the false gravid spot phenotype is expected to be lost in some *X. birchmanni* populations, suggesting that selection likely plays a role in its maintenance over time.

Balancing selection is expected to result in distinct patterns of variation in linked regions of the genome^66,67^, and we next investigated if the false gravid spot region contained such signatures. We again leveraged our higher coverage short-read dataset from the Coacuilco population, which had been collected agnostic to phenotype. Because we anticipated challenges with variant calling short-read data near the copy number variable regions within the structural variant, we focused on the 8.8 kb chromosomal inversion, which was present and single copy in all haplotypes identified with long-reads. Within the inversion, we found high nucleotide diversity (π), in the top 0.24% compared to 8.8 kb windows across chromosome 2 (Fig. 4e). Tajima’s D within the inversion was also somewhat elevated (top 6.97%), consistent with balancing selection (Fig. 4f). Finally, we calculated the within population non-central deviation statistic (NCD1) for the inversion, which has improved power compared to Tajima’s D to detect signatures associated with balancing selection^67,68^. We found that the inverted region was an outlier (bottom 0.97%), consistent with balancing selection acting on this region (Fig. 4g).

### Intra- and inter-sexual behavioral interactions provide clues about the mechanisms of selection on the false gravid spot

We next aimed to investigate mechanisms of selection that may favor or disfavor the false gravid spot polymorphism. Since male *Xiphophorus* aggressively guard access to food resources and mates^69–71^, we hypothesized that, by mimicking females, males with the false gravid spot may experience less aggression. Because the false gravid spot precedes development of other sexually dimorphic traits, we tested the effect of the false gravid spot in aggressive interactions during this life stage, when false gravid spot males might be expected to be more convincing female mimics. We designed experimental triads in the lab consisting of one large dominant male and two smaller size-matched males, one with and one without the false gravid spot. The smaller males were otherwise unornamented (i.e., lacked secondary sexual traits; Fig. 5a). Strikingly, we found that all dominant males (n=21) chased males with false gravid spot less than the paired stimulus male that lacked the false gravid spot (Fig. 5b; p<0.0001). Whether or not the large dominant focal male had false gravid spot did not significantly affect the frequency of aggressive interactions initiated (Fig. S19). Moreover, a scototaxis assay in *X. birchmanni* males raised in the laboratory yielded no differences in boldness between phenotypes (Fig. S20).

**Figure 5.**
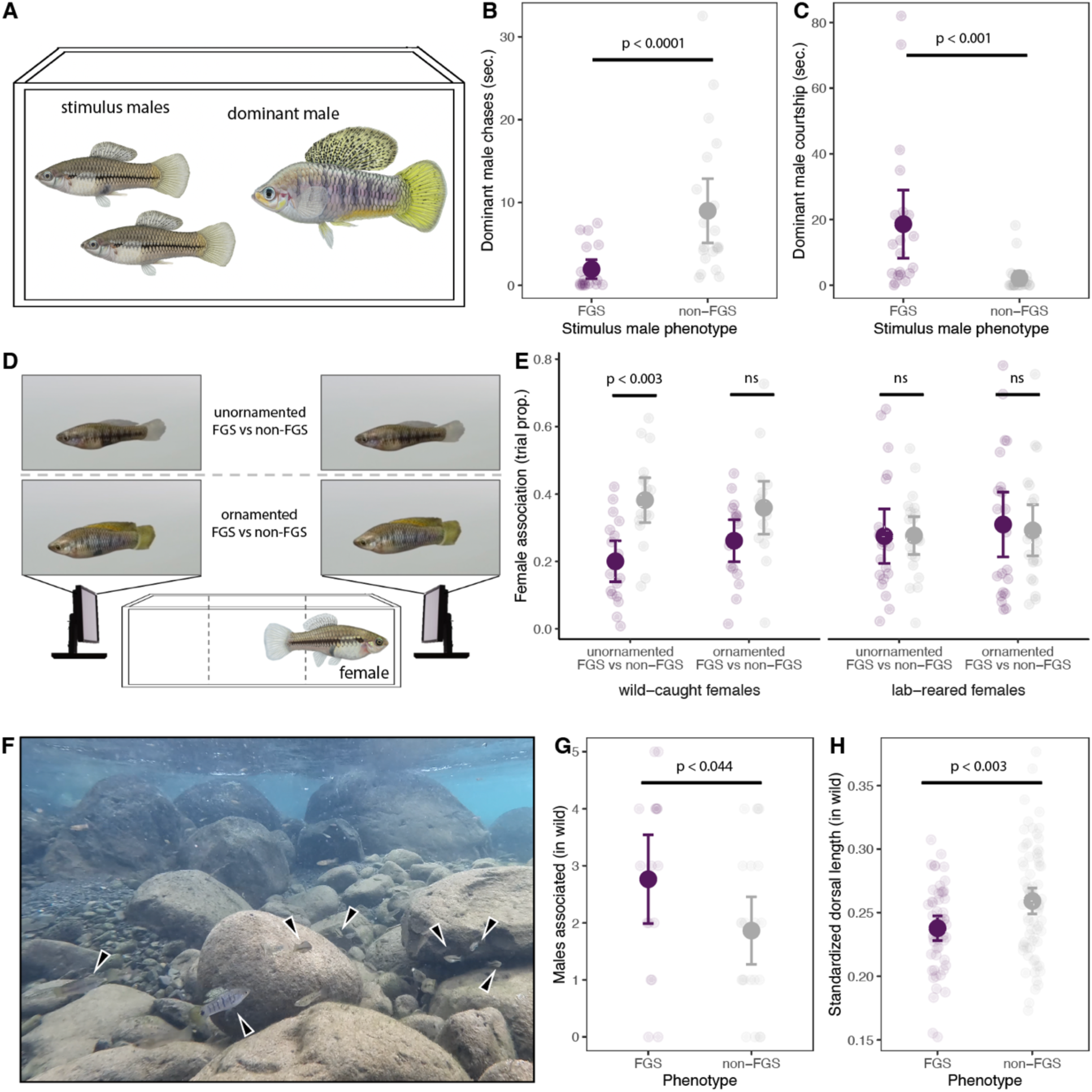
**Behavioral consequences of the false gravid spot. A**) Diagram of male-male interaction experimental design. Interactions initiated by the focal large-bodied, dominant male directed towards two smaller unornamented males, with and without false gravid spot were scored. **B**) False gravid spot (FGS) males are chased for less time compared to males without the false gravid spot (paired Wilcoxon test; p<0.0001). Large points represent experiment means ± 2 standard errors, with small points denoting mean value for each focal male. **C**) In the same trials, males with the false gravid spot were courted more than males without the false gravid spot (paired Wilcoxon test; p<0.001). **D**) Diagram of female preference experimental design. Females were presented a choice between false gravid spot and non-false gravid spot males in two animation types, unornamented males and ornamented males, with time spent in the 1/3 of tank closest to video stimulus considered association. **E**) Wild-collected females spend a greater proportion of the trial with animations of unornamented males without the false gravid spot (paired Wilcoxon test; p<0.003), suggesting disdain for the false gravid spot. A significant difference in preference was not detected in any other stimulus, or in a cohort of females born in the lab. **F**) Behavior observations in the wild were conducted in the Río Coacuilco with a mask and snorkel. Still image from a representative site during observations, with *X. birchmanni* noted with arrows. **G**) In the wild, FGS males were observed to be associated with more males in their immediate vicinity (1m^2^; GLM likelihood ratio χ^2^_1_ = 4.1, *p* < 0.044). **H**) Male *X. birchmanni* in the wild with false gravid spot have shorter dorsal fins than males without the false gravid spot (FGS; Welch’s t-test; p<0.002). Dorsal length is represented here as a fraction of body length.

While a reduction in aggression would likely provide a benefit to false gravid spot males, being perceived as female may lead to a cost if females experience greater harassment^31,72–74^. Because *X. birchmanni* court potential mates vigorously and males in other *Xiphophorus* species are known to attend to intensity of gravid spots when directing courtship towards females^43^, we also hypothesized males with the false gravid spot may receive additional sexual attention from the dominant male. Consistent with this, in our experimental triads, we observed that dominant males spent more time courting false gravid spot males than they did non-false gravid spot males (Fig. 5c). Taken together, our experimental assays demonstrate the false gravid spot is an important signal that other males attend to during this life stage.

Because female *Xiphophorus* (along with many species of poeciliids), choose mates based on a variety of characters^40,62^, including pigmentation^39,63,75^, we reasoned that the false gravid spot might be important for mate-choice dynamics. To assay preference, we ran dichotomous choice tests using video animations of male *X. birchmanni.* Because the false gravid spot develops before male-diagnostic sexually dimorphic traits, but remains expressed throughout the male’s life, it was unclear if female preference might depend on the presence of other male-diagnostic traits. We therefore tested female preference for males with and without the false gravid spot in two animation types (Fig. 5d): unornamented males (i.e., small dorsal fin, muted coloration; Video S1) and ornamented males (i.e., large dorsal fin and pigmentation; Video S2). Moreover, because decades of behavioral study in *Xiphophorus* and other poeciliids have demonstrated that female mate preference is dependent on experience, we simultaneously tested a cohort of wild-caught females and a cohort reared in the laboratory with no previous mate choice experience. Females from the wild associated 40% less with false gravid spot males in the unornamented phenotypic background (Fig. 5e; p=0.003), suggesting females disdain the false gravid spot in some contexts. We found a trend for female disdain of the false gravid spot in the ornamented males, but this was not significant (Fig. 5e; p=0.11). In contrast, we did not find evidence of preference for or against the false gravid spot in females that were born in the laboratory (Fig. 5e; p=0.89). Simulations revealed that we expect to have good power to detect large effect sizes in our experiments (e.g., ∼90% power based on the inferred effect size in our experiment focusing on unornamented males with wild-caught females) but low power to detect weaker effect sizes (e.g., 32% in our experiment focusing on ornamented males in the same group). This suggests that although we do not find significant disdain for the false gravid spot in trials with ornamented males, we cannot rule out the possibility that female disdain for the false gravid spot is merely weaker in this scenario (Fig. S21).

Based on insights from manipulative experiments in the laboratory, we next sought to evaluate impacts of the false gravid spot in its natural context. During behavioral observations of 51 individuals conducted at the Río Coacuilco (Fig. 5f), we found that inter- and intra-sexual social interactions were generally less frequent than in the laboratory experiment, with both male and female *X. birchmanni* spending most of their time swimming against the current or feeding. However, we observed males with the false gravid spot were associated with significantly more males in close proximity (Fig. 5g; GLM likelihood ratio χ^2^_1_ = 4.1, *p* < 0.044). Because the increased density of nearby males in the wild could be consistent with either the false gravid spot reducing male-male aggression or increasing male-male courtship, we searched for additional clues as to the condition of males with false gravid spot in natural populations. We did not detect differential growth rates of juveniles with and without false gravid spot in the wild (Supplementary Information). However, in other *Xiphophorus* species, the growth of secondary sexual ornaments depends on food availability and the amount of aggression males experience^48,76^. To begin to investigate this in *X. birchmanni*, we measured male dorsal fin length in wild-caught males, since dorsal fins are an important male ornament in *X. birchmanni*^62,77^. Intriguingly, we found the dorsal fin ornament was on average 10% smaller in *X. birchmanni* males with the false gravid spot (Fig. 5h, Welch’s t-test; p<0.003), which could be consistent with lower food acquisition or higher stress, although we did not directly observe differential feeding rates in the wild (p=0.24).

## Discussion

Phenotypic polymorphisms present opportunities to study scenarios in which natural selection may favor the maintenance of multiple alleles. To gain a comprehensive understanding of how such polymorphisms are controlled, we investigated the genetic and developmental basis of a sexual mimicry polymorphism in the swordtail fish *X. birchmanni* and explored behavioral mechanisms of selection that play a role in its maintenance. Together, this work contributes to an integrated understanding of the genetic mechanisms and evolutionary forces maintaining an understudied class of balanced polymorphisms.

Variation in color patterns were some of the first traits to provide insights into the inheritance patterns and frequency changes of adaptive phenotypes^2^ and remain tractable systems to investigate the genetic architecture and selective mechanisms acting on adaptive traits^78–80^. We identify structural variation 1.5 kb upstream of the *kitlga* 5’ UTR as the causal locus driving the false gravid spot phenotype in *X. birchmanni*. *kitlga* is related to the ancestral vertebrate *kitlg*, a well described pigmentation gene^50^, but was duplicated during the ancient teleost whole genome duplication (generating *kitlga* and *kitlgb*). In teleost fishes, *kitlga* has maintained the ancestral pigmentation function in multiple species^81,82^. *kitlg* and *kitlga* have been frequently implicated in pigmentation variation in natural populations, and previous work has indicated that they do so almost exclusively through *cis-*regulatory changes^83^, likely due to highly pleiotropic effects of protein-coding changes. Single-nucleotide changes in *kitlg* enhancer sequences that control pigmentation phenotypes have been identified in humans^84^. Differential *kitlga* regulation has been implicated in pigmentation variation in natural populations of stickleback^51^ and in domesticated populations of betta fish^85^, although the causal variants have remained elusive. However, tissue-specific regulation of ligands is thought to be a major driver of pigment pattern variation in fish by controlling the spatial distribution of chromatophores cells^86^. Our findings add to the emerging consensus that modification of similar genes and pathways underpin diverse pigmentation phenotypes^80,86^.

While the role of structural variants in the evolution of phenotypic polymorphisms is well appreciated in both theoretical and empirical work, much of the focus has been on large chromosomal inversions which may function as “supergenes”^16,17^. Interestingly, long-read sequencing in humans has revealed that smaller inversions (1-20 kb) are often associated with nearby structural variation, especially repeat expansions^87^. Smaller structural variants acting on single genes to generate phenotypic polymorphisms have been previously described^27,88,89^ but these have largely been interrogated with short-read data and coverage-based analyses, limiting their resolution and precision. Using long-read sequencing, we fully resolved complex structural variants that drive the false gravid spot phenotype. The 5 structurally variable haplotypes we identified among the 8 false gravid spot haplotypes sampled stand in contrast to the simple non-false gravid spot haplotype, where we detected only one structural variant (a singleton insertion) despite surveying a larger number of haplotypes (Fig. 2c). The high structural diversity of the false gravid spot haplotype lends support to the idea that structural variation can predispose sequences to the evolution of additional structural variation, driven by the complexities of DNA replication, repair, and recombination in such regions^87,90^.

The location of this structural variant in a non-coding region suggested that it may directly play a role in altering expression of neighboring genes. Together, our RNA-seq and allele-specific expression results demonstrate that the localization of the structurally complex haplotype upstream of *kitlga* results in both tissue-specific and allele-specific upregulation of *kitlga* in males with false gravid spot. While it remains unclear which of several structural differences within the false gravid spot haplotype might drive variation in expression, copy number variation in non-coding regions have been associated with eQTL and disease phenotypes^90,91^. Although the importance of these elements has not been studied in many species, our findings raise the possibility that smaller scale structural variation may be an important driver of phenotypic polymorphisms through its impact on gene regulation. Addressing this question is newly tractable for a diversity of species in the era of long-read sequencing and chromatin accessibility assays.

Several lines of phenotypic and genetic evidence show that balancing selection is acting to maintain the false gravid spot polymorphism in *X. birchmanni*. We found the false gravid spot affects social interactions with both males and females, which may provide insights into the mechanisms of selection acting upon this trait. One clear benefit of female mimicry is that males with the false gravid spot experience less frequent aggression from other males, reducing the risk of direct injury from aggressive interactions. Such aggressive interactions are common in male *X. birchmanni* and can result in mortality in the lab environment. However, we also identified two potential costs of the false gravid spot in both male-male and male-female interactions. Despite being chased less, males with false gravid spot were courted more by other males compared to males without the trait. While these interactions are not directly competitive, male harassment may be energetically costly and time spent being courted represents time that males may not be able to feed^31,73,74^. In the wild, we found that false gravid spot males also had smaller ornaments, which could be consistent with an energetic cost^48^. Additionally, females disdain males with the false gravid spot in some situations, which could drive differential mating success via female mate choice. Taken together the false gravid spot trait likely impacts fitness in natural populations.

While we see evidence for both costs and benefits of the false gravid spot in particular situations, how exactly they interact to maintain the trait in natural populations remains unclear. Theory suggests that opposing fitness effects alone can only maintain balanced polymorphisms under a narrow range of circumstances^92–94^. Our data may be consistent with several different forms of balancing selection including spatiotemporally varying selection and negative frequency dependent selection. In these scenarios, selection coefficients vary based on context, resulting in the maintenance of multiple phenotypes. While we did not test for this directly, our behavior experiments may hint at some context dependence. When shown animations of males without ornamentation, females displayed strong disdain for the false gravid spot, but in animations with ornaments, females showed a trend towards weaker discrimination against false gravid spot males (though this may be impacted by power). Moreover, strength of preference may also vary as a function of female experience. We speculate that context dependence in social interactions could allow the false gravid spot to be maintained by frequency-dependent selection or environmentally fluctuating selection, which are commonly involved in alternative mating systems^22,64^. Disentangling these possibilities and uncovering other selective mechanisms that impact this naturally occurring balanced polymorphism represent exciting directions for future work.

## Methods

### Sample collection

*X. birchmanni* samples used for this study were collected using baited minnow traps from sites across the Mexican states of Hidalgo, San Luis Potosí, and Veracruz, with permission from the Mexican government and Stanford University animal welfare protocols (Stanford APLAC protocol #33071). Samples from the genome-wide association study were collected from the Río Coacuilco (21°5’51.16″N 98°35’20.10″W) between 2017 and 2018 for a previous association mapping experiment^36^. Fish used to generate long-read assemblies were collected between 2021-2023 from Coacuilco, Benito Juarez (20°52’51.51″N 98°12’24.05″W), and Izapa (21°1’22.81″N 98°56’16.99″W). Fish used in gene expression analyses were born and raised in the Stanford fish facility between 2021-2023. Fish tested in behavior experiments were collected from Coacuilco using the same methods, except when otherwise noted. Fish used in the otolith experiment were collected in 2023 from Coacuilco. In most experiments that required lethal sampling, fish were euthanized with an overdose of MS-222 followed by severing of their spinal cord. However, in the gene expression experiments, we anesthetized fish on ice and euthanized fish by severing the spinal cord.

### Morphological analyses and histology

Male *X. birchmanni* were dissected under a dissection microscope to examine the structure of the false gravid spot. Two non-false gravid spot males and two false gravid spot males were dissected. Three anatomical components of the false gravid spot were identified. We also examined the structure of the false gravid spot using a histology-based approach. We fixed two male *X. birchmanni* with the false gravid spot, two without, and two female *X. birchmanni* in a 4% formaldehyde solution for 24 hours at 4 C. Following fixation, the tissue was dehydrated, embedded in paraffin and we generated transverse sections for imaging at 5μm thickness. These sections were stained with hematoxylin and counterstained with eosin.

### Sample phenotyping

To obtain phenotypes, fish were anesthetized in a buffered solution of MS-222 and water at a concentration of 100 mg/mL. Anesthetized males were photographed with their dorsal and caudal fins spread on a grid background and ruler for reference, using a Nikon d90 DSLR digital camera with a macro lens. Because lighting environment was not standardized for photographs taken in the field, we recorded binary phenotypes for presence or absence of false gravid spot. We took quantitative measurements of male standard length, body depth, dorsal fin length and height using Fiji^95^. We also recorded presence of melanic patterns (e.g. vertical bars, horizontal line, carbomaculatus, spotted caudal) and xantho-erythrophore coloration.

### Genome Wide Association Study

We reanalyzed previously published low-coverage (∼0.35×) whole-genome Illumina data from 329 male *X. birchmanni* collected between 2017 and 2018^36^. The false gravid spot segregated at intermediate frequency in this natural population (126 with and 203 without), and occurred at similar frequencies in adult (0.39) and juvenile (0.38) males. As a result, we included both juveniles and adults in the association mapping analysis. We mapped all reads to the *X. birchmanni* reference genome using bwa-mem with default parameters and removed alignments with a mapping quality score less than 30. We then conducted a case-control GWAS using samtools-legacy program^96^ (https://github.com/lh3/samtools-legacy/blob/master/samtools.1) to estimate allele frequency differences between cases and controls and used a likelihood ratio test to determine association of each SNP with false gravid spot presence or absence. Given the presence of structural variation in this region of the *X. birchmanni* genome, we also repeated the analysis using an alternate reference sequence, with chromosome 2 replaced with the *X. birchmanni* false gravid spot haplotype (see below).

To determine the appropriate significance threshold for our analysis, we used a permutation-based approach. We randomly shuffled phenotypes between individuals, re-ran the case-control GWAS analysis using the samtools-legacy program, as we had for the real data, and recorded the minimum p-value. We repeated this procedure until we had collected 500 replicates. We took the lower 5% quantile of this distribution of minimum p-values and used this value (3.7 x 10^-9^) as our genome-wide significance threshold.

### Correcting for population structure in Genome Wide Association Study

The case-control design we used in GWAS analysis can be susceptible to a high false positive rate in the presence of population substructure, because population substructure can also generate allele frequency differences between cases and controls, despite such allele frequency differences not specifically being driven by the trait being mapped. This is a particular concern with traits like the false gravid spot that may have a function important in mating (and thus be linked with assortative mating^97^).

Because the data we collected is low-coverage, standard approaches to control for population structure are not well-suited for our data. Nonetheless, we implemented several analyses to explore the potential effects of population structure on our results. We first generated “pseudo-haploid” calls for each individual by generating pileup files with bcftools^96,98^, randomly sampling a read from each position, and assigning the allele supported by that read as the allele present in that individual. If a site was not covered by any reads in an individual, it was coded as missing. After excluding sites with a minor allele frequency less than 2% or missing in 75% of individuals using plink^99^ --recode, we performed PCA analysis with plink using this data. We asked whether there was a correlation between false gravid spot phenotype and any of the first 10 PCs, which together explained ∼28% of the variation in the data.

We also used these pseudo-haploid calls to repeat the GWAS with plink while explicitly accounting for population structure by including the first 4 PCs as covariates. Despite the expectation that we would have reduced power due to pseudo-haploid calls, we still detected an association on chromosome 2 that surpassed the genome-wide significance threshold (Fig. S3). Since different iterations of an analysis using pseudo-haploid calls will differ due to stochasticity introduced by sampling, we repeated this procedure 10 times. We found that the p-value for the association between pseudo-haploid calls in this region of chromosome 2 and the false gravid spot trait was always less than 1x10^-10^.

### Long-read Xiphophorus reference assemblies

We generated a new chromosome-level *X. birchmanni* genome using PacBio long-read sequencing. The reference individual was selected from a lab-raised individual from stocks derived from the Coacuilco population and had no false gravid spot. Genomic DNA was isolated from tissue using the Qiagen Genomic-Tip kit. We followed the manufacturer’s protocol with slight modifications. Tissue was digested in 1.5 mL of Proteinase K and 19 mL of Buffer G2 for two hours. The sample was gently mixed by inversion every 30 minutes. The sample was applied to the column following equilibration of the column with 10 mL of Buffer QBT. The column was washed two times with 15 mL of Buffer QC, and then eluted in a clean tube with Buffer QF. DNA in the eluate was precipitated using 10.5 mL of isopropanol, mixed by gentle inversion, and then centrifuged at 4 C (5000 x g) for 15 minutes. The pellet was washed in cold 70% ethanol, re-pelleted, and then air dried after removal of the supernatant in 1.5 mL of Buffer EB. DNA was quantified on the Qubit fluorometer and evaluated for quality using the Nanodrop and Agilent 4150 TapeStation machine. Extracted DNA was sent to Cantata Genomic for PacBio library prep and sequencing on 2 SMRT cells. Reads were initially checked for quality using NanoPlot^100^ and residual adapters were removed using the HiFiAdapterFilt.sh script^101^. We used hifiasm^102^ with default parameters to generate a phased diploid genome assembly and scaffolded the primary contigs output using previously collected Hi-C data^36^ with YaHS^103^ to generate chromosome-level assemblies. All chromosomes were oriented and named with respect to the *X. maculatus* reference genome. Because chromosome 21 is thought to be the sex chromosome in most *Xiphophorus* species^49^, we used minimap2^104^ to aligned both alternate haplotypes generated by hifiasm to identify the alternate haplotype. We found a ∼2 Mb region of highly divergent and structurally variable sequence on the distal end of chromosome 21. To represent both haplotypes (X and Y) in the assembly, which has been shown to improve variant calling performance on the sex chromosomes^105^, we made a second copy of chromosome 21 with the divergent region replaced with the alternate haplotig. We then aligned these draft X and Y scaffolds together and masked the high homology (pseudo-autosomal) regions on the Y, as recommended^105^.

Because the *X. birchmanni* reference individual did not have false gravid spot, we sought to assemble a high-quality reference for the false gravid spot haplotype. To do so, we also sequenced one *X. malinche* × *X. birchmanni* F_1_ hybrid with false gravid spot with PacBio HiFi, with extraction methods identical to those described above. Given the false gravid spot is absent in *X. malinche*, the false gravid spot allele in the hybrid must be inherited from *X. birchmanni.* We chose to sequence an F_1_ hybrid because we anticipated the 0.5% divergence between *X. birchmanni* and *X. malinche* would lead to an assembly with long phased blocks. Extracted DNA was sent to Admera Health Services, South Plainfield, NJ for PacBio library prep and sequencing on 2 SMRT cells. We checked read quality and removed residual adapters as above and again used hifiasm to generate a phased diploid genome assembly. Based on our initial GWAS results, we blasted *kitlga* to the diploid assembly graph (untigs) to identify the *X. birchmanni* and *X. malinche* haplotypes corresponding to the candidate region. We ran the resulting 2 untigs that contained *kitlga* through the ancestryHMM pipeline^106^. One of the untigs, a 23 Mb sequence, was inferred to be 100% *X. birchmanni* ancestry based on the ancestryHMM pipeline, suggesting this block was fully phased for the majority of chromosome 2. We then manually replaced the homologous non-FGS haplotype in the reference with this untig based on coordinates generated using a minimap2 alignment, following the approach described above for chromosome 21.For consistency, we also reassembled previously published *X. malinche* HiFi reads^107^ using the computational pipeline described above and added previously collected Hi-C data^36^.

### Annotation of reference assemblies

We annotated the new reference assemblies for *X. birchmanni* following a pipeline used for previous *Xiphophorus* assemblies. We first identified repeats using RepeatModeler^108^. We took the output file from this process and repeat libraries from Repbase^109^ and FishTEDB^110^ were input into RepeatMasker^111^, allowing additional transposable element sequences to be identified based on sequence similarity. These annotations were used to hard mask transposable elements and soft-mask simple repeats in subsequent annotation of protein-coding genes.

We annotated protein coding genes using a multi-pronged approach that included homology, transcriptome mapping, and *ab initio* prediction. For annotation using homology, we collected a total of 455,817 protein sequences from the following databases: the vertebrate database of Swiss-Prot (https://www.uniprot.org/statistics/Swiss-Prot), RefSeq database (proteins with ID starting with “NP”’ from “vertebrate_other”) and the NCBI genome annotation of human (GCF_000001405.39_GRCh38), zebrafish (GCF_000002035.6), southern platyfish (GCF_002775205.1), medaka (GCF_002234675.1), mummichog (GCF_011125445.2), turquoise killifish (GCF_001465895.1) and guppy (GCF_000633615.1). We used GeneWise^112^ and exonerate to align these protein sequences to the *de novo* assembly to identify gene models via homology, using GenblastA^113^ to identify an approximate alignment region and improve required computational time.

We also took advantage of available RNA-seq data from multiple tissues from F_1_ hybrids between *X. birchmanni* and *X. malinche* to perform transcriptome mapping. The RNA-seq data was pre-processed using fastp^114^ and mapped to the *de novo* assembly using the program HISAT2^115^. We processed mapping results with StringTie^116^ to generate gene models based on this data. We also used Trinity^117^ to assemble *de novo* transcriptomes from the RNA-seq data and aligned these transcripts to the assembly. These aligned transcripts were converted to gene models using Splign^118^. Finally, we performed *ab initio* gene prediction using AUGUSTUS^119^. The first round of AUGUSTUS training was performed on BUSCO genes and the second round of training was performed using gene models identified from the methods descripted above. This database was used as ‘hints’ for AUGUSTUS gene prediction.

With all these predictions in hand, we generated a final consensus annotation by screening models by locus. When two models competed for a splice site, we retained the gene model that was better supported by transcriptome data. When a terminal exon (with start/stop codon) from *ab-initio* or homology gene model was better supported by transcriptome than that of the retained gene model, the latter was replaced. We also kept an *ab-initio* prediction when its transcriptome support was 100% and it had no homology prediction competing for splice sites.

Given that automatic annotation approaches, such as those described above, can miss exons separated by large introns and thus lead to incomplete gene models, we inspected the annotation of *kitlga*. In particular, visualization of RNA-seq data aligned to the *X. birchmanni* reference genome with HISAT2, and the gene models of *kitliga* for several tissues generated by stringtie, suggested an additional first exon and UTR 30 kb upstream of the second exon, and 1.5 kb downstream of the structural variant (Fig. S4). While this has implications for interpretation of the GWAS peak, for downstream analyses involving gene expression, we did not modify the transcriptome to maintain consistency with other genes in the genome (e.g. for RNA-seq analyses).

### Analyses of protein evolution at kitlga

We compared amino acid sequences at *kitlga* between *X. birchmanni* individuals with and without the false gravid spot to *X. malinche*, the sister species of *X. birchmanni* which is fixed for the non-false gravid spot phenotype. We reasoned that if coding changes were involved in phenotypic differences between individuals, we would see this reflected in this analysis of amino acid sequences. We extracted the predicted exons from previously generated pseudoreference sequences for 25 *X. birchmanni* individuals^120^ and five *X. malinche* individuals^121^. We found no segregating variation in the amino acid sequence at *kitlga* within these *X. birchmanni* individuals, nor within *X. malinche*. Moreover, *X. birchmanni* and *X. malinche* had identical predicted amino acid sequences (Fig. S6). However, they differed in several synonymous substitutions which we take advantage of below for allele specific expression analysis (Fig. S14). To calculate the rate of nonsynonymous substitutions per nonsynonymous site versus the rate of synonymous substitutions per synonymous site between *X. birchmanni* and *X. malinche* at *kitlga* we used the codeml program in PAML^122^.

### Linkage disequilibrium analysis in the region implicated by GWAS

To investigate haplotype structure in the associated region identified with GWAS, we used a previously collected dataset of 23 individuals from the Coacuilco population sequenced at higher coverage (median 18.4×). Reads were mapped to the *X. birchmanni* reference genome using bwa, and realigned around indels using picardtools. We called variants using GATK4^123^ and filtered each vcf individually following guidelines outlined in the GATK best practices that we previously verified have good performance in *Xiphophorus*^124^. We additionally filtered all SNPs within 5 bp of an indel. We then merged all SNP calls and removed sites with >10% missing data using bcftools. We used plink to calculate linkage disequilibrium (LD) for all SNPs within 1 Mb surrounding the associated SNP located in the center of the GWAS peak. We also genotyped individuals in this region based on sets of SNPs that are diagnostic for the rearranged haplotype, finding 2 homozygote false gravid spot, 8 heterozygotes, and 16 homozygous non-false gravid spot individuals.

### Long read sequencing and assembly of males with different false gravid spot phenotypes

For 11 of the 12 individuals collected from 3 populations as described above, we extracted high molecular weight DNA from brain and liver tissue with the NEB Monarch HMW DNA Extraction Kit for Tissue (Catalog #T3060S, NEB, Ipswich, Massachusetts). The purified DNA was size selected for >40kb fragments with the PacBio SRE XL buffer (Catalog #102-208-400, PacBio, Menlo Park, California). Long-read sequencing libraries were then generated by Oxford Nanopore R10.4.1 sequencing. Approximately 3 μg of size-selected DNA was prepared with the Oxford Nanopore ligation sequencing kit (Catalong #SQK-LSK114, Oxford Nanopore, Oxford, United Kingdom) following the manufacturer’s instructions, except we used half volumes of library prep reagents and extended bead elution steps from 5 minutes to several hours at 37 C to allow very long DNA fragments to resuspend completely. We loaded 200-300 ng of prepared library onto a R10.4.1 PromethION flow cell and sequenced on a P2 Solo sequencer with live basecalling with the fastest model enabled. After our targeted sequencing depth was achieved, the data were basecalled again with Guppy v6.4.6 (Oxford Nanopore) using the super-accuracy model. Reads passing the default quality score filter (Phred-scaled QV10) were assembled using the diploid assembler shasta^125^ with default parameters.

For the remaining individual, a male from Coacuilco, we extracted genomic DNA using the Promega Wizard HMW DNA extraction kit (Catalog #A2920, Madison, WI), following the manufacturer’s instructions. DNA was quantified on the Qubit fluorometer and evaluated for quality using the Nanodrop and Agilent 4150 TapeStation machine. Extracted DNA was sent to the University of Washington Long Reads Sequencing Center for PacBio library prep and sequencing on 1 SMRT cell. A diploid assembly was generated from the HiFi reads with hifiasm as described above.

### Analysis of structural variation at the false gravid spot locus

For an initial analysis of structural variation comparing the *X. birchmanni* non-false gravid spot reference, the *X. birchmanni* false gravid spot reference sequence, and the *X. malinche* assembly, we performed pairwise all-to-all alignments in MUMmer4^52^. For population level analyses from long-read assemblies, we first identified phased haplotypes containing the structural variant by conducting BLASTn^126^ searches for the inverted region against the diploid assemblies (Assembly-Phased.fasta for nanopore assemblies and *utg.fa for HiFi assemblies). We chose to use these files, which are produced as part of the assembly pipelines, because they contain all the sequence information but are more conservative regarding assembly errors (for example, untigs have fewer assembly errors than contigs^102^). We aligned these phased haplotypes to the *X. birchmanni* non-false gravid spot reference genome using MUMmer4 with default parameters and visualized the results using a custom R script. For ease of comparison between haplotypes, we also used MUMmer4 to align regions identified in the reference alignment (segmental duplications, inversion, transposable element insertions) to each phased assembly. We visualized the regions of homology using the gggenes R package (https://cran.r-project.org/web/packages/gggenes), keeping segments with ³1 kb length and merging consecutive segments separated by less than 500 bp.

We initially tested several approaches to quantitatively call structural variation at the focal locus on chromosome 2 from long-read data. Due to the complex nature of the rearrangement associated with the false gravid spot, we were limited to structural variant callers that recognize both inversions and insertions/deletions (indels). We tried variant callers that take advantage of data from both reads (e.g. Sniffles2^127^ and SVision^128^) and assemblies (PAV^87^). While these approaches always identified structural variants in false gravid spot haplotypes, the results were not consistent in the number and nature of variants called. Given this inconsistency, we report results of the MUMmer-based analysis of structural rearrangements in this region.

In addition to pairwise alignments, we also constructed a local pangenome of the region using the pan genome graph builder (PGGB)^54^ with parameters -s 500 -p 90 -k 35. We trimmed the graph with ODGI^129^ and visualized it in Bandage^130^. Homology within the graph to each region identified in MUMmer4 was determined using BLASTn searches natively implemented in Bandage.

### Evaluation of DNA structural features near GWAS peak

We used available computational tools to compare the frequency of non-B DNA in the regions surrounding the false gravid spot and non-false gravid spot haplotypes using the program nBMST (https://github.com/abcsFrederick/non-B_gfa). This program outputs the coordinates of predicted features associated with non-B DNA for each fasta sequence analyzed, including A-phased repeats, G-quadraplexes, Z-DNA, direct, inverted, and mirror repeats, and STRS. We compared both the frequency and the length of these features between the phased false gravid spot and non-false gravid spot reference haplotypes, locally around the GWAS peaks identified mapping to both reference sequences.

### Whole tissue RNA-seq

To explore expression differences of *kitlga*, as well as other genes, in individuals with and without the false gravid spot, we dissected 3 tissues from 13 individuals: brain, body wall musculature, and the tissue surrounding the gonopodial suspensorium which consists of the erector analis major muscle and its perimysium. We extracted total RNA with the Qiagen RNAeasy Mini Kit (Catalog #74106, Qiagen, Valencia, CA) and sent RNA to Admera Health Services for library preparation using the NEBNext Ultra II Directional library prep kit with Poly A Selection. Extracted RNA was sent to Admera Health Services, South Plainfield, NJ for library preparation and sequencing on Illumina HiSeq 4000. Extractions and library preps were paired between groups (e.g. at least one false gravid and non-false gravid replicate for each extraction and library prep batch) to control for batch effects in statistical analysis downstream.

Raw reads were trimmed to remove adapter sequences and low-quality base pairs using cutadapt and TrimGalore! with parameters --phred33 --quality 30 -e 0.1 --stringency 1 --length 32 --paired --retain_unpaired^132^. Trimmed reads were pseudoaligned to the *X. birchmanni* reference transcriptome with *kallisto*^56^ and the differential expression analysis was performed using DESeq2^57^. Because the *X. birchmanni* reference transcriptome does not contain isoforms, we assumed one transcript per gene id in the reference genome. We removed genes with zero counts in all samples before beginning analysis of differential expression with DESeq2. We analyzed each of the three tissue types separately and used an analysis model that modeled gene expression as a function of false gravid spot phenotype and extraction/library preparation batch. Gene counts were normalized by library size and we used the “local” function in DESeq2 to estimate within-group dispersion. We tested for significant differences in expression using the Wald test and calculated shrunken log-fold changes in expression using the ashr package^133^. We consider genes with an adjusted p-value<0.05 between groups to be differentially expressed.

### rt-qPCR quantification of kitlga expression

As a secondary confirmation that *kitlga* was differentially expressed in the pigmented erector analis major and its perimysium, we took a real-time quantitative PCR approach. We extracted total RNA using the Qiagen RNAeasy Mini Kit (Catalog #74106, Qiagen, Valencia, CA) following the manufacturer’s instructions, and used the GoScript Reverse Transcription System kit to generate cDNA from extracted RNA (Catalog #A5000, Promega Corporation, Madison, WI). We designed primers using the tool primer-Blast^134^ and the predicted cDNA sequence of *kitlga* (Fig. S14; Table S3). We tested the specificity of these primers using amplification of cDNA with the Phusion PCR kit (Catalog #M0530, Thermo Scientific, Wilmington, DE) and visualizing the amplicon size on an Agilent 4200 Tapestation (Agilent, Santa Clara, CA). We next quantified amplification efficiency for rt-qPCR. We generated a serial dilution series of *X. birchmanni* cDNA and ran these samples in triplicate on a BioRad CFX384 C1000 touch real-time thermocycler machine (Biorad, Hercules, CA) with no-template negative controls. The per-reaction mastermix was made up of: 1 μl cDNA, 0.3 μl of the forward and reverse qPCR primers, 5 μl of Maxima SYBR Green mastermix (Catalog #K0253, ThermoFisher Scientific, Waltham, MA) and nuclease free water. The cycling conditions used were 95 C for 10 minutes, followed by 40 cycles of 95 C for 30 seconds, 62 C for 1 minute, and a read step. Each run included a final melt curve and were inspected for evidence of a single PCR product. *efa1* was used as a housekeeping gene. Both selected primer sets had efficiency between 108-112%.

### Allele specific expression of kitlga in lab-generated hybrids heterozygous for the false gravid allele at kitlga

Based on our RNA sequencing results, we wanted to determine if differential expression of the *kitlga* allele associated with the false gravid haplotype was driven by changes in *cis*. Because there was little sequence variation within *X. birchmanni* in the *kitlga* coding region, but several fixed synonymous differences between *X. birchmanni* and its sister species *X. malinche* (Fig. S14), we conducted this experiment using hybrids between the two species. Because *X. birchmanni* is polymorphic for the false gravid spot whereas *X. malinche* is fixed for its absence, we used early generation hybrids (F_2_-F_3_) and selected individuals with the false gravid spot. We confirmed by genotyping that these individuals were heterozygous for *X. birchmanni* and *X. malinche* ancestry in the region surrounding *kitlga* using the AncestryHMM^106^ pipeline.

We designed primer sets for pyrosequencing using the Qiagen Pyromark software. We tested several primer sets on three tissues (brain, gill, testis) derived from pure *X. birchmanni* and pure *X. malinche* cDNA. Briefly, we amplified samples (N=3 of each species) using the PyroMark PCR kit (Catalog #978703, Qiagen, Germantown, MD) following the manufacturer’s instructions. PCR reactions were submitted to the Protein and Nucleic Acid facility (Stanford University, CA) for pyrosequencing. We analyzed our results using the Qiagen Pyromark software. Based on the results of these quality tests, we selected three primers that performed well (>95% support for the expected allele in all individuals of both species; Table S3). We repeated this approach to collect data from 7 hybrids with the false gravid spot in 4 tissues: brain, gill, testis, and erector analis major and its perimysium. We calculated the ratio of the false gravid spot (*X. birchmanni*) allele to the non-false gravid spot allele (*X. malinche*) and tested if this ratio differed from 1 using a Welch’s t-test.

### Developmental timing of false gravid spot in X. birchmanni

To determine the developmental stage at which the false gravid spot develops, we tracked individuals throughout development. We established two 115 L tanks of newborn *X. birchmanni* fry from lab stocks. We expected these fish to be segregating for the false gravid spot based on the observed phenotypes in the adult stocks. We set up tanks with ∼25 fry and maintained them following normal husbandry procedures until they reached 4-5 cm of length. Once fish reached this size, they were marked with a unique color combination using elastomer injection tags (Northwest Marine Technologies). Fish were checked weekly for evidence of gonopodial differentiation. As soon as males could be unambiguously determined based on thickening of the anterior segments of anal fin ray 3, they were separated into a grow-out tank and photographed weekly. During these photographs, we also examined males under the dissection scope and scored the stage of gonopodial development (Fig. S16), ranging from thickening of the 3^rd^ ray (onset of external phenotypic sexual differentiation) to a fully mature gonopodium capable of fertilization with differentiated distal structures including a claw, blade, hook, spines and serrae (Fig. S16; Table S4). Phenotypic data was collected from these weekly photographs as described above.

To understand if the presence of the false gravid spot influenced the development of other secondary sexual traits, including morphological and pigmentation traits, we ran a principal components analysis on adult males in this dataset using the prcomp function in R. We considered measurements of standard length, body depth, longest dorsal ray, and dorsal length as morphological variables. For pigmentation, we considered binary presence versus absence of xanthophore pigmentation in the dorsal and caudal fin, melanic pigmentation in the vertical bars, horizontal line, spotted caudal, and carbomaculatus. X. birchmanni *population divergence, demography, and simulations of likelihood of maintenance of polymorphisms*

To better understand the demographic history of *X. birchmanni* populations and divergence between them, we mapped nanopore reads to the reference *X. birchmanni* assembly using minimap2 -x mapont and called variants using longshot^135^, a variant caller designed for single-molecule reads with higher error rates than Illumina reads, with default parameters. Using these variant calls, we generated “pseudoreference” sequences in the same coordinate space following established workflows in the lab and used custom lab scripts to calculate pairwise sequence divergence (dxy) between each individual (https://openwetware.org/wiki/Schumer_lab:_Commonly_used_workflows; https://github.com/Schumerlab/Lab_shared_scripts). We used the R package pheatmap to cluster the results (Fig. S18). To infer demographic history for each population, we next ran PSMC^64^ using longshot variants for each sample. To scale the output, we used 3.5 x 10^-9^ as the mutation rate and 0.5 years for generation time, following previous work^124^.To explore the likelihood of maintenance of a neutral polymorphism in simulated populations matching the inferred demographic histories, we performed forward-time population simulations with SLiM^65^. We tracked evolution at a single site, prevented recurrent mutations at that site, and directed the program to track substitutions as well as polymorphisms. In generation 2, we used the genomes.addNewDrawnMutation function to establish a polymorphism at the tracked site at the expected allele frequency based on observed phenotype frequencies in each population (Benito Juarez: 0.13, Izapa: 0.23, Coacuilco: 0.29). We took the estimated N_e_ for each time segment from PSMC from a single individual from each population. Then, we implemented population size changes at time segment intervals output by PSMC using setSubpopulationSize to make the simulated population exactly match the inferred N_e_ at that timepoint. Due to differences in population history, we lose resolution in demographic inference at different time periods. For Izapa, which has undergone a strong bottleneck, we lose resolution in PSMC inference ∼65k generations in the past. Although PSMC inference for Coacuilco and Benito Juarez extends to deeper timescales, we selected the starting time segment for these populations that most closely matched where we lose resolution in Izapa to begin simulations for each population. This ranged from 63-73k generations in the past for Coacuilco and Benito Juarez. After initializing the simulation, we allowed evolution to occur until the focal polymorphism fixed or was lost or until the simulation ran to completion. We recorded how frequently the polymorphism was retained in each population across 10,000 replicate simulations for each population.

### Detecting genomic signatures of balancing selection

To pursue independent lines of evidence that the false gravid spot is maintained by balancing selection, we next investigated patterns of genomic diversity near the false gravid spot locus. Because only a small number of individuals were sequenced with long-read approaches, and because combining data across populations can generate artifacts in these analyses due to population structure, we again used previously generated short-read data generated for 23 individuals from the Coacuilco population. An additional benefit of using these individuals is they were collected randomly with respect to false gravid spot phenotype, and therefore represent a less biased sample of the population. Due to potential technical issues with mapping and variant calling short-read data near and within the copy number variable regions within the structural variant, we focused on the 8.8 kb chromosomal inversion. This region distinguishes false gravid spot and wild-type individuals but was present and single-copy in all haplotypes analyzed with long-reads, averting issues with mappability generated by copy number variable regions. Using variant calls generated as described above, we first calculated mean nucleotide diversity (π) using pixy^136^ in 8.8 kb windows across chromosome 2. We next used vcftools^137^ to calculate Tajima’s D within the inversion and compared it to windows of the same length chromosome-wide. Lastly, we used the R package balselr (https://github.com/bitarellolab/balselr) to calculate NCD1 statistics in windows chromosome-wide of the same size. For this analysis, we used a target frequency of 0.3, based on the empirical observation of allele-frequency at the FGS locus, but found that in practice the NCD1 results were not sensitive to using different target frequencies.

### Frequencies across and within *X. birchmanni* populations

To evaluate variation in the frequency of the false gravid spot in *X. birchmanni*, we visited 9 populations across the species range. We phenotyped adult males collected using baited minnow traps for presence or absence of the false gravid spot. To understand how much false gravid spot frequency varies over time, we also sampled phenotypic frequencies from the Coacuilco *X. birchmanni* population from 2017-2023. Again, we phenotyped only adult male fish for the false gravid spot. We asked whether the frequency of the false gravid spot significantly differed across years in these populations using a binomial test.

### Male-male interaction open arena behavioral experiments and analysis

To test whether the presence of false gravid spot affects male-male behavioral interactions, we set up triads consisting of one large focal male (n=20), and two sized matched small target males–one with false gravid spot and one without–in 40L (50.8 x 25.4 x 30.5 cm) open-field arenas. All target males lacked sexually dimorphic traits that typically develop in males later in life. All arenas were equipped with two half terracotta pots for shelter and all sides of each arena except the front and top were covered on the exterior with black plastic to achieve visual occlusion between trial arenas. The interactions of each focal male were video recorded from the front for 15 minutes (GoPro Hero10 camera on a tripod) twice daily for 4 consecutive days. Trials ran for five weeks (Tuesday-Friday) using 4 arenas total. The day before the trial, a dominant male and female were added to a new tank to habituate for at least 16 hours. Each day the first 15-minute video was recorded in the morning within 5 minutes after introducing the two size-matched, randomly selected stimulus males to the trial arena. The second video was recorded five to six hours later in the afternoon. Following the afternoon recording, stimulus males were removed from the tank and, and each focal male was moved to a new arena with a new female to habituate overnight before the next trial. This procedure was followed to ensure that no focal male saw the same targets twice and no target male was used more than once in each week. All fish were housed in single sex colonies for at least two weeks prior to testing. All holding tanks and test arenas were illuminated from above on at 12/12hr light dark cycle. All tanks were fed normally once daily between video recordings.

The first five minutes of each 15-minute video was discarded as the acclimation period and behaviors were scored for the remainder of each video. This resulted in a total of 80 minutes of observation per focal male across trials. The following behaviors were scored from recorded videos with the aid of computer assisted event logging software (BORIS^138^ v.7.13.9): Number of chases – rapid movement by focal male toward a target resulting in the target rapidly swimming away; number and duration of sustained chases – chases where focal male continued to pursue target after initial interaction; number and duration of courtship bouts – characterized by side by side dorsal flaring and circling of target by focal male; number and duration of aggressive displays – characterized by dorsal flaring while facing in the opposite direction (head to tail) often accompanied by tail beating and nipping behaviors; gonopodial flexes – the rotating forward of the gonopodium, a behavior associated with ejaculation preceding mating attempts in poeciliid fish; time spent hiding; number of approaches – when the focal male swims to within one body length of a target individual without chasing or engaging in either courtship or aggressive displays. Total time spent chasing was calculated by multiplying the total number of non-sustained chases by 0.5 seconds, the typical length of such interactions, and adding the duration of sustained chases. All measures except hiding were categorized by the target to which the focal male was directing its attention.

Unless otherwise stated, for statistical analyses, means across all videos recording the focal male were used (i.e. the four triads of which a given focal male was a member; 8 observations total per focal male). Because most experiments yielded a non-normal data distribution, we used non-parametric Wilcoxon signed rank tests for all analyses comparing interactions of focal males with the pair of stimulus males with and without false gravid spot.

### Dichotomous trial behavioral experiments and analysis

To test if females displayed a preference or disdain towards the false gravid spot, we ran a series of dichotomous choice tests using animations of male *X. birchmanni.* Females originated from two lab stocks from the Coacuilco population: a wild-caught cohort (n=17) and a cohort born in the laboratory (n=23). Each trial was run for 55 minutes with a 10 minute acclimation period. The animations (Fig. 5d) were of two types (1) paired unornamented males with and without false gravid spot and (2) paired ornamented males with and without false gravid spot. Animations were generated using models based on images of adult males without the false gravid spot, and a false gravid spot was computationally added to the relevant animation. Animations were made using blender and scaled to be the same standard length (based on the average adult standard length in the Coacuilco population). To control for side bias, after a 5 minute rest period we displayed identical animations with the location of the false gravid spot male on the opposite monitor. Each trial animation was 5 minutes long, and in total each female saw 4 videos (differing in ornamentation and side of the stimulus). The order in which fish saw trials was random, except side-bias trials were shown sequentially, meaning a total of 4 possible trial orders were done. The rest and acclimation periods showed a screen with the tank background, but no fish animation stimulus.

We used EthoVisionXT16^139^ to track movement of fish during the female preference behavioral trials. Due to low grayscale contrast between the fish and the bottom of the tank, we used color marker tracking. We chose a marker color range for subject identification that was as narrow as possible while still picking up fish in all areas of the tank. We created arena settings that encompassed the full tank as much as possible, while excluding anything that fell into the marker color range, such as shelters, the edges of the tank, and small reflections on the water. We created two preference zones on each side of the tank, one 25cm from the video screen, and a closer nested zone 5 centers from the video screen. The center of the tank was the neutral zone. After tracking acquisition at a sample rate of 7.73 samples per second, we used EthoVision’s track editor feature to manually correct the location of the tracked fish as needed. This primarily occurred when the fish was on the very edge of the tank or within the shelter. We defined the start of a trial after the female visited both zones. We analyzed the difference in proportion of the trial spent in association with either stimulus. Because this experiment yielded a non-normal data distribution, we used Wilcoxon signed rank tests to compare differences in the proportion of time with the null expectation of 0.

### Scototaxis trial behavioral experiments and analysis

To study if the false gravid spot influenced boldness, we performed a scototaxis assay in males born in the laboratory. Fish typically prefer a dark tank background and time spent on a white background is a routinely used measure of boldness^140,141^. We used the males that were analyzed for the male development tracking study after data collection for that experiment had concluded. We constructed trial lanes (l x w x h; 50 cm x 19 cm x 19 cm) and then lined one half with white and one half with black custom cut matte pvc foam board to reduce reflectivity. We rinsed the trial lanes between experiments with fish water and performed a water change. We recorded videos from above for a trial duration of 5 minutes and used EthoVisionXT16^139^ to track movement of fish during the scototaxis trials. Time spent in the white zone and latency to enter the white zone were scored as a measure of boldness. To control for side bias, we tested each fish twice with the zones flipped and took the mean value of the amount of time spent on the white or black background across the two trials. We kept the standard ethovision experiment settings mode and contour-based center-point detections for this set of trials. We chose the differencing detection mode with a fixed background image and the setting for the subject darker than the background, which accurately tracked the fish when in the white zone. Because the fish was not reliably detectable when in the dark zone, we designated that half of the tank as a “hidden” zone in the Arena settings. Whenever the fish entered the hidden zone, its position was assumed to be at the center of the zone. We created a tracking zone in the white zone but left an approximately 1 cm gap between the edge of the hidden zone and the white zone so that fish were not counted as entering the light zone until their entire body had left the hidden zone. After tracking acquisition at a rate of 6.25 samples per second, we used EthoVision’s track editor feature to manually correct the location of the tracked fish as needed. Because this experiment yielded a non-normal data distribution, we used non-parametric Wilcoxon signed rank tests to compare males with and without false gravid spot in time spent in the white zone and in latency to enter the white zone.

### Field observations and statistical analyses

Using snorkel and masks, we conducted focal observations in the Río Coacuilco. After selecting an area where there was at least one *X. birchmanni* male, we waited for at least 3 min for the fish to habituate to our presence. After the habituation period we counted the numbers of males (identified by the presence of a gonopodium) and females (identified by the presence of a gravid spot and absence of a gonopodium), within one square meter of the focal male. We scored males for the presence of the false gravid spot, and as covariates, we also scored the focal male for body size and size of dorsal fin. Because we were unable to directly measure the fish, we scored size as either smaller or larger than 44 mm reference line, based on the average male standard length for this population. We also recorded dorsal fins as either “small” or “large” for their body size. We observed males for 5 minutes or until the male swam out of the area, remaining at least 1 m away from the focal male when possible. Each observer switched between watching a male with and without false gravid spot in the same location. In addition, each observer moved to a new area in the river between these pairs of observations so as not to observe the same males more than once. The behaviors we recorded were the following: number of courtship displays performed by the focal male toward a female, number of gonopodial thrusts (i.e., attempted copulations), number of times the focal male chased and did aggressive displays to another male (i.e.; aggressive behaviors); and number of times they were chased (i.e., retreats). In addition, we recorded the number of nips at the substrate (feeding).

We performed all statistical analyses using Generalized Linear Models (GLMs) with a normal probability distribution and an identity link function. After building initial models that included all the independent variables, we performed a model reduction using Akaike’s information criterion corrected for small sample size (AICc), while retaining the variables with significant effects (assessed by their *P* values) and until we found a minimal model. In cases where the normality assumption of residuals was not fulfilled, we used a different probability distribution and link function until a normal distribution of residuals was confirmed. We used two independent GLMs to assess the effects of the morphological traits we scored for focal males on the number of males and females around them (dependent variables). For the analyses of the focal male interactions with both males and females, we grouped the behaviors according to their function (aggression and courtship). To simplify analyses, we used a single measure of the grouped behaviors (first principal component; PC1) calculated from factor analyses that included chases and displays to males (aggression) and displays and attempted copulations to females (courtship). In these cases, the full models included the three morphological traits and their interactions as factors, but also the number of males around (in the GLM of aggression behaviors) and females around (in the GLM of courtship behaviors) as covariates.

### Morphological correlates of the false gravid spot in the wild

To understand if there were any morphological correlates of the false gravid spot in wild fish, we measured fish used in the previously collected GWAS population between 2017 and 2018. We measured male standard length as well as dorsal fin length and width using Fiji^95^. We used linear models to assess if these traits were affected by false gravid spot phenotype and sample year. We initially fit a full model containing both false gravid spot phenotype and sample year, but in our final analysis we dropped sample year since we did not find that it was correlated with morphology (p>0.05).

## Acknowledgments

We are grateful to the Federal Government of Mexico for permission to collect fish. We thank members of the Schumer lab, Peter Sudmant, Jeff Groh, and Rishi De-Kayne for helpful comments on earlier versions of this manuscript. We thank Dorian Noel for providing fish illustrations and Dylan Keim for creating the blender animations. Stanford University and the Stanford Research Computing Center provided computational support for this project. TOD and NBH were supported by NSF GRFP awards DGE-2146755, BMM was supported by NSF GRFP award DGE-1656518, and SMA was supported by a Stanford Science Fellowship. This research was funded by the Searle and Pew Foundations and by a Freeman Hrabowski award to MS, and a Stanford Tinker award to MS and CGR. This research was also supported by funding from the National Science Foundation (IBN 9983561) and Ohio University (Research Incentive) to MRM.

## Data availability

Data will be made available on Dryad and NCBI SRA. Code is available at https://github.com/tododge/xbirchmanni_fgs/ and https://github.com/Schumerlab/.

## Author contributions

Conceptualization, T.O.D., D.L.P., and M.S.; Investigation, T.O.D., B.Y.K., J.J.B., S.M.B., T.R.G., A.E.D., L.A.G., A.R.R., S.K.H.C., M.L.W., R.C., B.M.M., K.H., N.B.H., J.A.M.C., H.R.G., M.S.T, C.G.R., O.R.C., M.R.M., M.Scha., D.L.P., and M.S.; Software, T.O.D., K.D., D.L.P., and M.S.; Formal Analysis, T.O.D., L.A.G., O.R.C, M.R.M, D.L.P, and M.S.; Resources, M.S.; Writing – Original Draft, T.O.D., D.L.P, and M.S.; Writing – Review & Editing, T.O.D., B.M.M., S.M.A, O.R.C., M.Scha., D.L.P, and M.S.; Supervision, S.M.A., M.Scha, D.L.P, and M.S., Funding Acquisition, M.R.M and M.S.

