## Supplementary Information for "Complex structural variation and behavioral interactions underpin a balanced sexual mimicry polymorphism"

### Supplementary methods, results, and figures

*Reference assemblies for* X. birchmanni *and* X. malinche

We generated a new chromosome-level reference genome assembly for *X. birchmanni* using PacBio HiFi and Hi-C data from a male without false gravid spot. We also re-assembled a reference genome for the sister species *X. malinche*[1], a species in which the false gravid spot polymorphism is absent. The resulting assemblies were highly contiguous and complete, with contig N50s of 26.1 Mb for *X. birchmanni* and 10.7 Mb for *X. malinche* and BUSCO completeness scores of 98.7% for both species. Following integration of previously collected Hi-C data[2], scaffold N50s were 32.3-32.6 for *X. malinche* and *X. birchmanni* respectively. The 24 largest scaffolds contained 98.7% and 98.4% of the assembled sequence for *X. birchmanni* and *X. malinche*, respectively, matching the 24 chromosome karyotype typical of *Xiphophorus* species.

To generate the alternate false gravid spot haplotype for *X. birchmanni*, we assembled a phased genome assembly from an $F_1$ hybrid between *X. malinche* x *X. birchmanni* with the false gravid spot. Due to mendelian inheritance, this $F_1$ hybrid was heterozygous for the false gravid spot and necessarily inherited the false gravid haplotype from *X. birchmanni* (the father in this cross design) because the false gravid spot is absent from *X. malinche.* We chose to assemble a genome from an $F_1$ hybrid because we expected that the divergence of *X. malinche* and *X. birchmanni* (0.5% per bp) would facilitate generating long phased blocks (N50 of untigs: 12.3 Mb), which would prove useful for examining structural differences within and between species. Using a BLASTn search we identified 2 PacBio "untigs" that contained the *kitlga* coding sequence in this $F_1$ assembly, and ran both through the ancestryHMM pipeline[3]. Both untigs were inferred to derive 100% of their ancestry either for *X. birchmanni* or *X. malinche,* suggesting that this $F_1$ assembly approach generated blocks that were fully phased. We selected the 23 Mb untig that was 100% *X. birchmanni* ancestry as the false gravid spot haplotype.

*Identifying diagnostic false gravid spot SNPs between* X. birchmanni *haplotypes*

To identify SNPs diagnostic of false gravid spot haplotypes, we leveraged the long-read dataset collected across populations. For simplicity, we focused on the region that is inverted in false gravid spot haplotypes, since this region single-copy in all haplotypes. To identify diagnostic SNPs in the region, we first mapped all haplotypes to the *X. birchmanni* non-false gravid spot reference using minimap2. We then used IGV to search the 8.8 kb inverted region for single nucleotide differences between haplotypes that were fixed between the inverted and non-inverted haplotypes. We excluded SNPs that were not fixed in either haplotype as well as SNPs within 5 bp of an indel. Together, this analysis identified 147 fixed differences between haplotypes that we use as "diagnostic SNPs" in analyses presented in the main text.

*Power analyses of dichotomous mate choice trials*

Dichotomous choice trials are intended to measure the strength and direction of female preference for a given signal. In practice, differences detected in preferences for different stimuli or between different groups of females in behavioral trials could reflect true differences in the strength of preference, or instead point to differences in power between experiments. To explore these possibilities, we performed simulations to determine our power to detect preferences of various effect sizes given our experimental design and sample size. We calculated difference in preference for each individual and then determined the mean, standard deviation, and sample size for each of the 4 groups. These groups were: 1) lab-born females tested against ornamented stimuli, 2) lab-born females tested against unornamented stimuli, 3) wild-caught females tested against ornamented stimuli, and 4) wild-caught females tested against unornamented stimuli. For each scenario, we then generated simulated data for each group using a range of effect sizes for preference (from 0 to 0.4 in increments of 0.025). Specifically, here effect size refers to the difference in the proportion of time a female spends with the two stimuli. As an example, to generate an effect size of 0.4, a hypothetical female could spend 60% of the trial associated with the non-false gravid animation, 20% with the false gravid animation, and 20% with neither stimulus. To generate the simulated data, we assumed a normal distribution with the mean set to the focal effect size and used the observed standard deviation from the trial of interest and the sample size from the trial of interest. In each simulated dataset, we determined whether the estimated effect size differed from zero using a Wilcoxon test at $p<0.05$, as we had in the analysis of the empirical data. To calculate expected power for each effect size and for each group, we calculated the proportion of tests where we detected an effect out of 10,000 replicate simulations.

This analysis revealed that we expect to have similar power for 3 of the 4 experiments (Fig. S21), with lab-born females tested against ornamented stimuli experiment having lower expected power than the other groups. As expected, our power increased when effect sizes were larger, with power exceeding 80% power in most groups at an effect size of ~0.175. This suggests that for experiments with large effect sizes, such as those we observe in trials of wild-caught females tested against unornamented stimuli, we have excellent power to detect preference. However, at somewhat weaker effect sizes, we expect to have poor power to detect an effect of interest. For example, we predict only ~35% power to detect an effect size of ~0.1. We note that this effect size is similar to the estimated empirical effect size from the experiment testing wild-caught females tested against ornamented stimuli.

*Behavioral observations in natural populations*

While lab-based behavioral studies allow precise measurements and reduced variability due to controlled environmental and social conditions, we are ultimately interested in how the false gravid spot impacts behavior in natural populations. To this end, we conducted behavioral observations in the field (Methods). While false gravid spot was the primary focus of the study, we also scored other male traits, particularly their dorsal size, since this is an important ornamentation phenotype in *X. birchmanni*, as well as male body size, and report these results below.

False gravid spot did not appear to influence access to females as measured by females in the vicinity of males (Fig. S22). Instead, we found that the size of male's dorsal fin was the trait that best explained access to females (GLM: Likelihood ratio $\chi^2_1 = 4.9$, P = 0.027). Males with larger dorsal fins were typically found in association with more females (Fig. S23). Past work has suggested that, female mate preference for the dorsal fin size appears is highly variable, with some females preferring larger dorsal fins and some preferring smaller [4,5]. Feeding rates were primarily influenced by male size (GLM: Likelihood ratio $\chi^2_1 = 4.4$, P = 0.036), regardless of dorsal fin size or number of males in the vicinity (both excluded from the minimal model).

Larger males had a higher feeding rate (Fig. S24); while males with or without false gravid spot did not differ in feeding rate (Fig. S25, GLM: Likelihood ratio $\chi^2_1 = 2.3$, $P = 0.132$). We did not find that any of the morphological traits scored could explain retreats (how often a male was chased); however, these results suggest that smaller males may be chased more often than larger males.

*Growth rates of juvenile males with and without the false gravid spot*

We wanted to investigate if growth rates differed as a function of false gravid spot phenotype during juvenile development in natural populations. To determine growth rates from wild-caught males from the Coacuilco population, we obtained ring counts from otoliths, the structures present in the inner ears of *X. birchmanni* males. Otoliths rings in juvenile *Xiphophorus* are deposited daily and are visually distinguishable from previously deposited rings under a microscope until males reach sexual maturity[137], at which point the otolith growth is dramatically reduced. Thus, we reasoned that otolith ring counts from juveniles could allow us to estimate the approximate age of each fish and thus calculate their growth rates. Otoliths from 48 wild-caught juvenile males with and without false gravid spot from Coacuilco were collected and preserved in 95% ethanol. The otoliths from both the right and left side of the head were removed and mounted to a microscope slide using permount (FisherScientific). The asteriscus (middle-sized otolith) was examined because this is the most translucent of the three otoliths. The otoliths were then imaged using a Nikon ECLIPSE Ti Series confocal microscope. At 40x magnification, we manually focused on the mid-plane of the otolith and took a Z-stack of 51 images at 0.5 micron slices. We then post-processed the image stacks two ways using Fiji. First, we compiled a max projection of each image stack and sharpened once and enhanced contrast by 0.35% (Fig. S26a). Second, we selected a single image from each stack and sharpened it once (Fig. S26b). To improve accuracy and reduce subjectivity, four researchers manually counted the rings of each otolith, using the cell counter plug-in in Fiji to track the rings using a color-coordinated system. Each of the 4 observers sequentially counted rings, moving the cell count marks until consensus was achieved (see Methods). Because the two image types produced similar counts with a weaker correlation than expected (Fig. S27; $R^2=0.52$), we took the mean of two counts for each otolith as an age estimate in days.

As expected, we found that otolith ring count (age) was significantly associated with the variation in standard length (Fig. S28; $p<0.0008$). However, we did not observe differences in growth rate in adult males with and without the false gravid spot (Fig. S29; $p<0.28$). This is perhaps not unexpected given adult male body size in the wild did not differ as a function of false gravid spot phenotype in our larger sample collected at Coacuilco from 2017-2018 (Fig. S30).

**Supplementary Tables**

**Table S1.** Results of DESeq2 analysis of differential expression between false gravid and non-false gravid males across several tissues.

| Gene id | log2FoldChange | p-value | symbol | tissue |
|---|---|---|---|---|
| g22533 | 3.726 | 7.61E-45 | PMEL | PM+EAM |
| g5347 | 4.104 | 2.68E-42 | tyrp1 | PM+EAM |
| g15929 | 3.267 | 2.71E-34 | TYRP1 | PM+EAM |
| g9067 | 3.276 | 5.66E-31 | MLANA | PM+EAM |
| g17743 | 3.072 | 1.03E-29 | mreg | PM+EAM |
| g13517 | 2.712 | 6.66E-21 | SLC2A11 | PM+EAM |
| g10491 | 2.897 | 3.98E-18 | Oca2 | PM+EAM |
| **g2346** | **1.993** | **1.24E-12** | **kitlga** | **PM+EAM** |
| g15639 | 3.876 | 2.30E-12 | - | PM+EAM |
| g20179 | 2.785 | 2.30E-12 | tyr | PM+EAM |
| g17988 | 2.253 | 9.06E-12 | Tspan10 | PM+EAM |
| g16685 | 2.197 | 1.24E-10 | CKB | PM+EAM |
| g10565 | 1.752 | 3.31E-10 | - | PM+EAM |
| g4660 | 2.479 | 9.26E-10 | slc24a5 | PM+EAM |
| g25700 | 2.002 | 3.08E-08 | ZDHHC2 | PM+EAM |
| g10070 | 2.265 | 7.78E-07 | ATCAY | PM+EAM |
| g23055 | 2.23 | 4.19E-06 | Kif5b | PM+EAM |
| g14511 | 1.673 | 7.58E-06 | slc30a8 | PM+EAM |
| g1977 | 1.634 | 8.81E-06 | DHFR | PM+EAM |
| g12495 | 1.724 | 1.24E-05 | tyr | PM+EAM |
| g10340 | 0.696 | 1.40E-05 | PRDX6 | PM+EAM |
| g18049 | 4.135 | 1.74E-05 | cyp2k1 | PM+EAM |
| g4237 | 4.488 | 1.87E-05 | Prss2 | PM+EAM |
| g7785 | 1.871 | 1.87E-05 | Fgf14 | PM+EAM |
| g9181 | 2.234 | 4.69E-05 | Slc45a2 | PM+EAM |
| g13239 | 1.955 | 5.79E-05 | adra2b | PM+EAM |
| g18048 | 2.86 | 7.02E-05 | cyp2k4 | PM+EAM |
| g15620 | 1.273 | 1.50E-04 | Fcgr1 | PM+EAM |
| g17347 | 1.544 | 1.50E-04 | OPN5 | PM+EAM |
| g24450 | 1.821 | 2.07E-04 | Hmx2 | PM+EAM |
| g24576 | 1.141 | 2.78E-04 | Arhgef33 | PM+EAM |
| g16256 | 2.239 | 3.17E-04 | - | PM+EAM |
| g25605 | 1.604 | 6.37E-04 | MCHR2 | PM+EAM |
| g13516 | 1.795 | 6.82E-04 | SLC2A11 | PM+EAM |
| g25329 | 2.117 | 8.28E-04 | SLC43A3 | PM+EAM |
| g13112 | 1.093 | 8.97E-04 | Rab27b | PM+EAM |

| | | | | |
|---|---|---|---|---|
| g26355 | 0.917 | 1.41E-03 | MLPH | PM+EAM |
| g16159 | 3.408 | 1.57E-03 | Cdon | PM+EAM |
| g20326 | 1.35 | 2.58E-03 | SLC30A1 | PM+EAM |
| g26404 | 0.342 | 3.24E-03 | DYNLT3 | PM+EAM |
| g1653 | 0.511 | 3.67E-03 | FAM180B | PM+EAM |
| g14087 | 0.335 | 3.83E-03 | C1orf216 | PM+EAM |
| g20182 | 1.546 | 3.83E-03 | Rab38 | PM+EAM |
| g9068 | 0.641 | 3.98E-03 | KIAA2026 | PM+EAM |
| g8175 | -0.619 | 4.24E-03 | MXRA5 | PM+EAM |
| g25419 | 1.609 | 4.68E-03 | BSCL2 | PM+EAM |
| g18014 | 0.534 | 4.97E-03 | lgals3bpb | PM+EAM |
| g8003 | 1.823 | 6.20E-03 | DCT | PM+EAM |
| g18018 | 2.548 | 7.08E-03 | - | PM+EAM |
| g812 | 1.747 | 7.17E-03 | DTNBP1 | PM+EAM |
| g21055 | 2.186 | 7.89E-03 | SLC10A1 | PM+EAM |
| g17804 | 0.844 | 8.93E-03 | Bax | PM+EAM |
| g6856 | -0.864 | 8.93E-03 | Znf521 | PM+EAM |
| g1706 | 1.171 | 1.01E-02 | Trpm1 | PM+EAM |
| g14199 | 0.916 | 1.21E-02 | - | PM+EAM |
| g15623 | 1.942 | 1.21E-02 | CD22 | PM+EAM |
| g1094 | 0.246 | 1.21E-02 | ada | PM+EAM |
| g26406 | 0.735 | 1.38E-02 | GHR | PM+EAM |
| g17068 | 0.911 | 1.39E-02 | Pde7b | PM+EAM |
| g14210 | -1.967 | 1.52E-02 | Mr1 | PM+EAM |
| g5820 | 1.123 | 1.71E-02 | - | PM+EAM |
| g17171 | -1.286 | 2.09E-02 | HMGXB3 | PM+EAM |
| g2876 | -1.286 | 2.09E-02 | HMGXB3 | PM+EAM |
| g6019 | 1.005 | 2.09E-02 | - | PM+EAM |
| g15560 | -1.092 | 2.17E-02 | GIMAP7 | PM+EAM |
| g10805 | -1.654 | 2.18E-02 | SYT2 | PM+EAM |
| g898 | 0.325 | 2.18E-02 | Cd300ld3 | PM+EAM |
| g23276 | -0.254 | 2.43E-02 | EMILIN2 | PM+EAM |
| g10425 | -1.419 | 2.67E-02 | - | PM+EAM |
| g22864 | 1.201 | 2.67E-02 | LDLRAD2 | PM+EAM |
| g1330 | 0.268 | 2.86E-02 | - | PM+EAM |
| g3380 | 0.442 | 3.13E-02 | C4 | PM+EAM |
| g18578 | 0.121 | 3.27E-02 | MYH9 | PM+EAM |
| g18249 | 1.302 | 3.61E-02 | Ighv1-72 | PM+EAM |
| g2290 | -0.489 | 4.38E-02 | MBL2 | PM+EAM |
| g16427 | -5.174 | 3.99E-27 | BTNL8 | Flank |
| g22533 | 2.01 | 5.02E-17 | PMEL | Flank |

| | | | | |
|---|---|---|---|---|
| g16159 | 5.846 | 5.50E-13 | Cdon | Flank |
| g5347 | 2.111 | 2.42E-10 | tyrp1 | Flank |
| g13970 | 7.017 | 4.41E-09 | Glipr2 | Flank |
| g15981 | -22.472 | 3.69E-08 | GIMAP7 | Flank |
| g17743 | 1.515 | 3.69E-08 | mreg | Flank |
| g15929 | 1.392 | 1.46E-07 | TYRP1 | Flank |
| g15639 | 3.625 | 2.83E-07 | - | Flank |
| g26369 | -0.99 | 3.65E-07 | - | Flank |
| g18478 | 7.35 | 3.76E-07 | Gvin1 | Flank |
| g9067 | 1.515 | 9.70E-07 | MLANA | Flank |
| g10491 | 1.966 | 1.11E-06 | Oca2 | Flank |
| g16251 | 20.028 | 1.20E-06 | - | Flank |
| g16354 | -1.865 | 1.20E-06 | Gimd1 | Flank |
| g2955 | -2.854 | 1.67E-06 | - | Flank |
| g16265 | -4.376 | 1.86E-06 | MFAP4 | Flank |
| g14213 | 17.927 | 3.24E-05 | - | Flank |
| g16101 | -3.535 | 5.90E-05 | - | Flank |
| **g2346** | **1.29** | **1.74E-04** | **kitlga** | **Flank** |
| g23808 | 5.573 | 1.09E-03 | Nceh1 | Flank |
| g16185 | 4.088 | 1.15E-03 | - | Flank |
| g15287 | 1.224 | 1.47E-03 | FCGBP | Flank |
| g17595 | 2.471 | 1.60E-03 | Samd14 | Flank |
| g24893 | -0.978 | 1.69E-03 | Tor1b | Flank |
| g16685 | 1.276 | 1.74E-03 | CKB | Flank |
| g20326 | 1.604 | 1.74E-03 | SLC30A1 | Flank |
| g11399 | 3.773 | 2.00E-03 | Hcar2 | Flank |
| g1074 | -2.069 | 2.31E-03 | G0s2 | Flank |
| g10889 | 1.324 | 3.08E-03 | Tecta | Flank |
| g15807 | -1.235 | 3.62E-03 | EIF5A | Flank |
| g19739 | 4.62 | 4.47E-03 | - | Flank |
| g9580 | 5.118 | 4.58E-03 | - | Flank |
| g14210 | -5.24 | 4.74E-03 | Mr1 | Flank |
| g17438 | -4.315 | 4.90E-03 | - | Flank |
| g9546 | 1.785 | 4.90E-03 | - | Flank |
| g12428 | -1.849 | 4.92E-03 | dgat2 | Flank |
| g20362 | 1.734 | 5.30E-03 | caspa | Flank |
| g16412 | -1.751 | 5.89E-03 | cxadr | Flank |
| g17486 | 1.137 | 6.23E-03 | krt13 | Flank |
| g17636 | 0.963 | 6.41E-03 | Mybpc2 | Flank |
| g5454 | -0.989 | 6.86E-03 | - | Flank |
| g13148 | 1.421 | 6.87E-03 | - | Flank |

| | | | | |
|---|---|---|---|---|
| g2703 | 1.537 | 9.92E-03 | rhbg | Flank |
| g10787 | -1.755 | 1.12E-02 | - | Flank |
| g481 | -0.843 | 1.32E-02 | hsp70 | Flank |
| g12451 | 2.523 | 1.38E-02 | MMP13 | Flank |
| g15001 | -1.106 | 1.38E-02 | - | Flank |
| g19896 | -1.225 | 1.38E-02 | RBP2 | Flank |
| g2533 | 2.142 | 1.38E-02 | Cpne4 | Flank |
| g25244 | -3.024 | 1.38E-02 | PCDHGA10 | Flank |
| g16269 | 2.859 | 1.73E-02 | - | Flank |
| g9068 | 0.387 | 1.75E-02 | KIAA2026 | Flank |
| g15575 | 1.175 | 1.77E-02 | TMPRSS15 | Flank |
| g2411 | 0.77 | 1.77E-02 | Mlkl | Flank |
| g8585 | 1.045 | 2.00E-02 | - | Flank |
| g21271 | 1.189 | 2.15E-02 | - | Flank |
| g4237 | 2.358 | 2.15E-02 | Prss2 | Flank |
| g12758 | 0.634 | 2.45E-02 | HPGD | Flank |
| g4660 | 1.194 | 2.55E-02 | slc24a5 | Flank |
| g19273 | -1.99 | 2.62E-02 | IAPP | Flank |
| g3402 | 0.37 | 2.62E-02 | St14 | Flank |
| g8003 | 1.096 | 2.62E-02 | DCT | Flank |
| g17988 | 0.812 | 2.70E-02 | Tspan10 | Flank |
| g7225 | 0.795 | 2.98E-02 | CFP | Flank |
| g17773 | -1.819 | 3.00E-02 | Cavin1 | Flank |
| g19669 | 0.699 | 3.16E-02 | LGALS4 | Flank |
| g7270 | 1.492 | 3.16E-02 | - | Flank |
| g2324 | -1.035 | 3.18E-02 | - | Flank |
| g10524 | 1.57 | 3.21E-02 | - | Flank |
| g15722 | 1 | 3.21E-02 | - | Flank |
| g16243 | 1.482 | 3.21E-02 | - | Flank |
| g24895 | -0.861 | 3.21E-02 | EGFL7 | Flank |
| g25929 | 0.647 | 3.21E-02 | CD200 | Flank |
| g9860 | -0.61 | 3.21E-02 | RGS1 | Flank |
| g11286 | 0.555 | 3.71E-02 | CALCOCO2 | Flank |
| g26404 | 0.328 | 3.82E-02 | DYNLT3 | Flank |
| g13517 | 0.804 | 4.06E-02 | SLC2A11 | Flank |
| g3380 | 0.806 | 4.06E-02 | C4 | Flank |
| g17569 | 0.619 | 4.10E-02 | Syngr1 | Flank |
| g12090 | 0.508 | 4.55E-02 | amer2 | Flank |
| g21707 | 1.168 | 4.58E-02 | - | Flank |
| g12569 | 0.913 | 4.62E-02 | Vwa7 | Flank |
| g15671 | 0.621 | 4.63E-02 | Cxcl3 | Flank |

| | | | | |
|---|---|---|---|---|
| g5313 | 0.902 | 4.91E-02 | - | Flank |
| g18018 | 3.295 | 3.54E-08 | - | Brain |
| g23218 | -5.439 | 1.71E-07 | Plcxd2 | Brain |
| g15639 | 1.875 | 7.60E-07 | - | Brain |
| g6314 | 0.383 | 1.37E-04 | - | Brain |
| g7164 | 0.338 | 6.22E-04 | PNP | Brain |
| g22141 | -3.525 | 6.54E-04 | - | Brain |
| g2290 | -1.24 | 7.80E-04 | MBL2 | Brain |
| g6827 | 0.485 | 2.16E-03 | tf | Brain |
| g2091 | 0.322 | 2.53E-03 | Lamb1 | Brain |
| g4681 | 0.32 | 3.78E-03 | Mfge8 | Brain |
| g11984 | 0.323 | 4.48E-03 | slc22a6 | Brain |
| g9930 | 0.459 | 4.48E-03 | Slc5a5 | Brain |
| g2312 | -0.371 | 4.66E-03 | Snupn | Brain |
| g12401 | 0.274 | 8.95E-03 | Trim16 | Brain |
| g4490 | 0.771 | 8.97E-03 | CILP | Brain |
| g14076 | -0.58 | 9.50E-03 | MAP7D1 | Brain |
| g3638 | 0.311 | 1.24E-02 | Fxyd3 | Brain |
| g24021 | 0.376 | 1.36E-02 | - | Brain |
| g24663 | 1.735 | 1.42E-02 | CDH1 | Brain |
| g15633 | 0.491 | 1.66E-02 | SIGLEC1 | Brain |
| g12789 | -0.413 | 2.09E-02 | THAP9 | Brain |
| g16159 | 1.243 | 2.10E-02 | Cdon | Brain |
| g6936 | 0.244 | 2.23E-02 | Higd1a | Brain |
| g20934 | 0.817 | 2.26E-02 | TY3B-G | Brain |
| g7711 | 0.202 | 2.31E-02 | LONRF2 | Brain |
| g113 | 0.214 | 2.70E-02 | - | Brain |
| g2551 | 0.73 | 2.79E-02 | TG | Brain |
| g7165 | 0.304 | 2.79E-02 | KCNJ10 | Brain |
| g9980 | -0.292 | 2.79E-02 | ATP13A3 | Brain |
| g21989 | 0.291 | 2.83E-02 | - | Brain |
| g2955 | -0.437 | 3.45E-02 | - | Brain |
| g19988 | 0.249 | 3.50E-02 | apoa1 | Brain |
| g7684 | 0.208 | 4.12E-02 | COX17 | Brain |
| g17650 | 0.552 | 4.55E-02 | CLDN4 | Brain |
| g15006 | 0.248 | 4.89E-02 | Slc6a19 | Brain |
| g23840 | -0.26 | 4.89E-02 | - | Brain |
| g15777 | 0.252 | 4.89E-02 | Plac8l1 | Brain |
| g16101 | -0.474 | 4.99E-02 | - | Brain |

130

**Table S2.** Primers used in rt-qPCR quantification of *kitlga* expression.

| Primer target | Forward | Reverse | Efficiency |
|---|---|---|---|
| *kitlga* | TCAGCTTCTTCGCCAAGTCC | ACCTTCCACACAACCAGGAAG | 108 |
| *efa1* (housekeeping) | CCCCTAACCTGACCACTGAA | GTGGGTCGTTCTTGCTGTCT | 112 |

**Table S3.** Primers used in pyrosequencing quantification of allele specific expression of *kitlga*, designed with the Qiagen Pyromark software. Performance in pure parentals refers to the estimated allelic support for the species-specific allele in two *X. birchmanni* and two *X. malinche* individuals.

| **Primer target** | **Forward primer** | **Reverse primer** | **Sequencing primer** | **Performance in pure parentals** |
|---|---|---|---|---|
| *Kitlga* site 57 | TGGAGAAGAAGAAGGCTTTACAG | O-GAGGTTCAGCGCATTTTTTT | TCTGATGGCACCTCG | 97-99% |
| *Kitlga* site 759 | O-TCCATTTCCTGCTGTTCATGAC | TTTTCCAGTTGCAGCTGAATGTAC | GCTGAATGTACCCCA | 98-99% |

**Table S4.** Criteria used to classify gonopodial differentiation into stages 1-5 for male developmental staging experiment. Based on Kallman and Schreibman[6].

| Gonopodial Stage | Criteria |
|---|---|
| Stage 1 (onset of external puberty phenotype) | • Ray 3 thickens<br>• Fin has definite acute angle at cephalodistal corner<br>• Rays 3,4,5 short<br>• Ray 4 longer than 3 and 5<br>• Rays 3-4-5 form a fleshy protuberance at end of fin |
| Stage 2 | • Rays 3-4-5 elongate beyond margin of fin<br>• Segments of ray 3 increase from 9 to 22 |
| Stage 3 | • Proximal spines differentiate on ray 3 (closer to body) |
| Stage 4 | • Distal serrae form on back half of ray 4 |
| Stage 5 (sexually mature) | • Blade appears at end of ray 3<br>• Gonopodium stiffens<br>• Skin shrinks, ends of spines flush with periphery<br>• Can see tips of proximal spines |

### Supplementary Figures

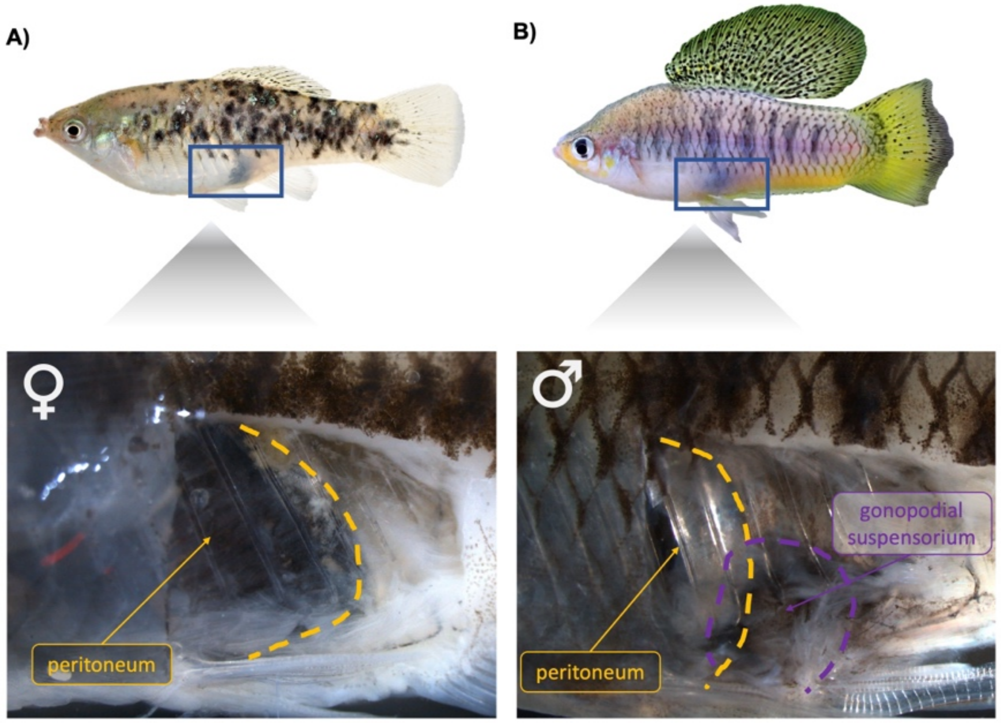


**Fig. S1.** Dissections of a female (A) and male (B) *X. birchmanni* reveal the gravid spot is a distinct structure from the false gravid spot. The gravid spot in females is generated by the pigmentation of the peritoneum, whereas the false gravid spot in males occurs posterior to the peritoneum and is due to pigmentation of the tissue surrounding the gonopodial suspensorium.

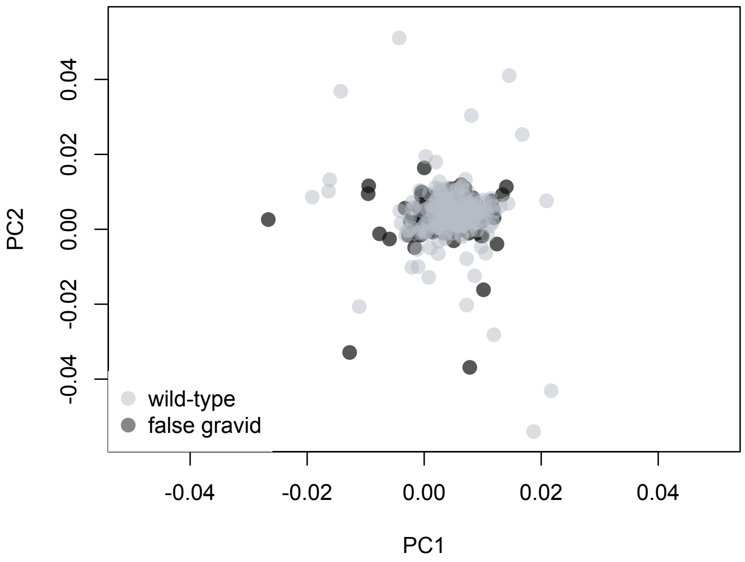


**Fig. S2.** We found no significant association between the false gravid spot and any of the first 10 principal components in a principal component analysis of genome-wide SNP variation. Plotted here are PC1 and PC2 with false gravid phenotypes highlighted (non-false gravid spot –light gray, false gravid spot – dark gray). We tested for correlations between phenotype and PC for the first ten PCs (see main text).

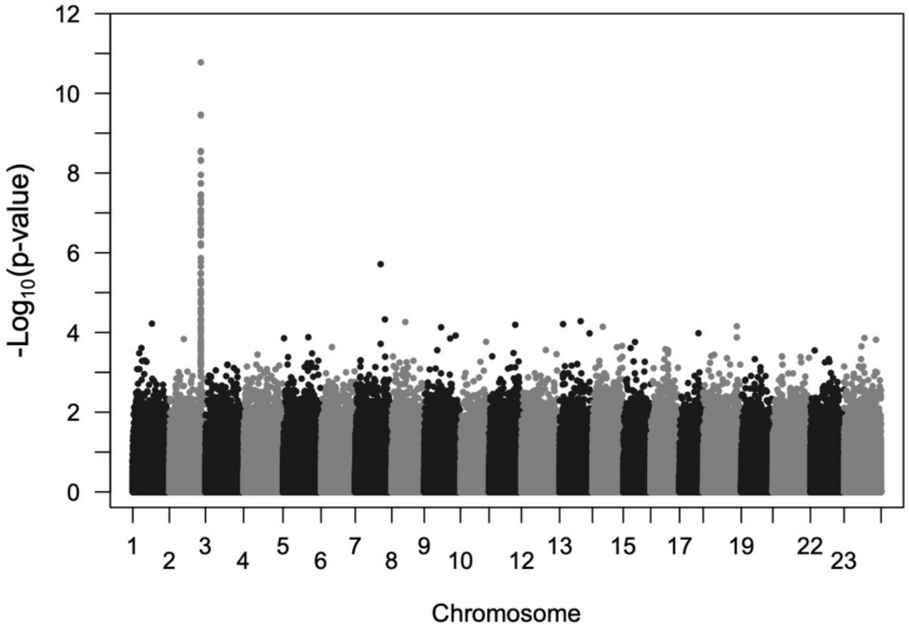


**Fig. S3.** Manhattan plot showing the results of a genome-wide association study of false gravid spot phenotype, accounting for PCs 1-4 as covariates in the analysis using plink. Although we have lower power in this analysis due to the use of pseudohaploid calls (see main text), we still detect the strong signal on chromosome 2 upstream of *kitlga*.

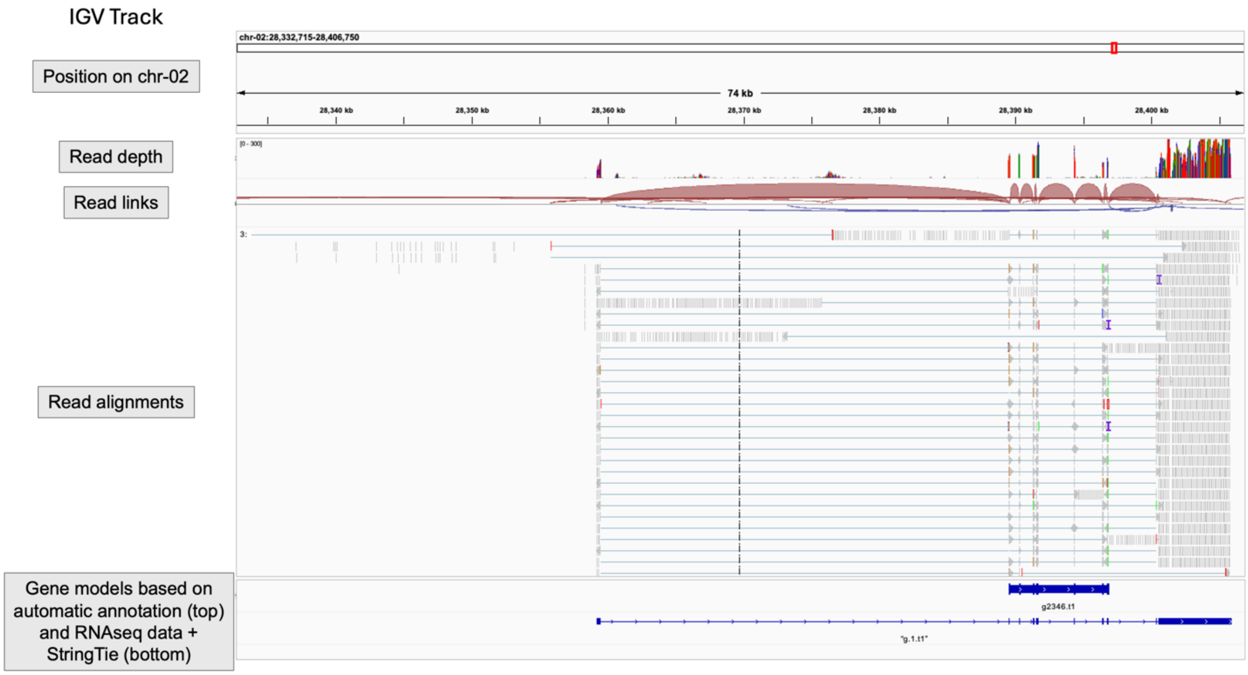


**Fig. S4.** Identification of *kitlga* exons based on RNAseq data mapped to the *X. birchmanni* reference genome with HISAT2. IGV snapshot showing alignment of RNAseq data from an $F_1$ individual that was used to annotate the reference genome. Top track shows the position of chromosome 2. The second track shows read depth, with the first putative exon achieving >100x read coverage per base-pair. The third track shows links between reads. The fourth track shows individual RNAseq reads mapped to the reference, with split alignments connected by blue lines. The fifth track shows results of the automatic annotation pipeline described in main text (top) as well as the results of our manual annotation described in the main text, based on the alignment and supported by gff files generated by StringTie (bottom). Note that an additional *kitlga* exon is supported by the RNAseq data and is included in our manual annotation.

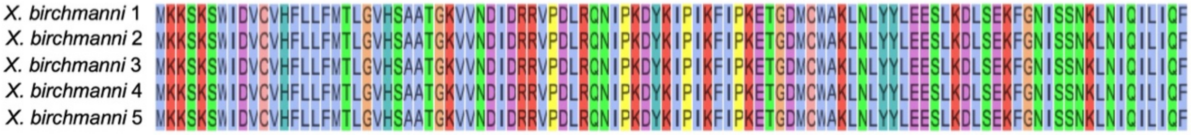


**Fig. S5.** Alignment of *kitlga* amino acid sequences from all available *X. birchmanni* pseudoreferences with complete sequence information using clustal-omega. Individuals with masked or missing basepairs in the coding sequence of *kitlga* were excluded from visualization here but were included in the analysis of amino acid sequence similarity across individuals.

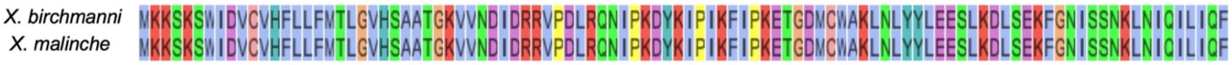


**Fig. S6.** Alignment of *kitlga* amino acid sequences from *X. birchmanni* and *X. malinche* reference sequences generated using clustal-omega. No nonsynonymous differences in *kitlga* sequence are detected between the two species.

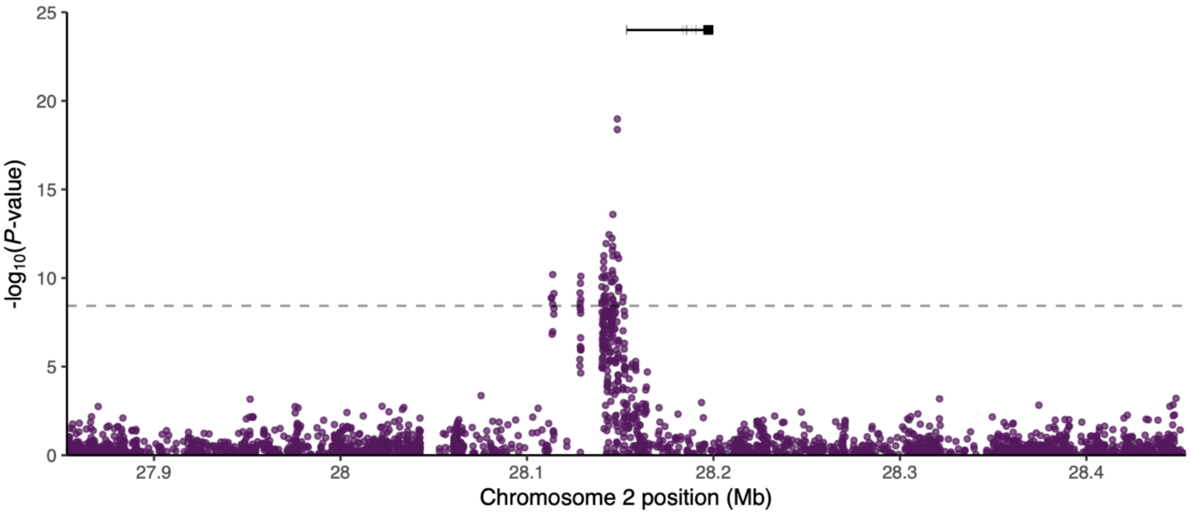


**Fig. S7.** Inset of GWAS results using the false gravid spot haplotype for chromosome 2. Gaps in the Manhattan plot correspond to the segmental duplications in this region, where Illumina short-reads do not map uniquely and are therefore filtered out of our analysis. The *kitlga* gene model is also shown above the GWAS peak.

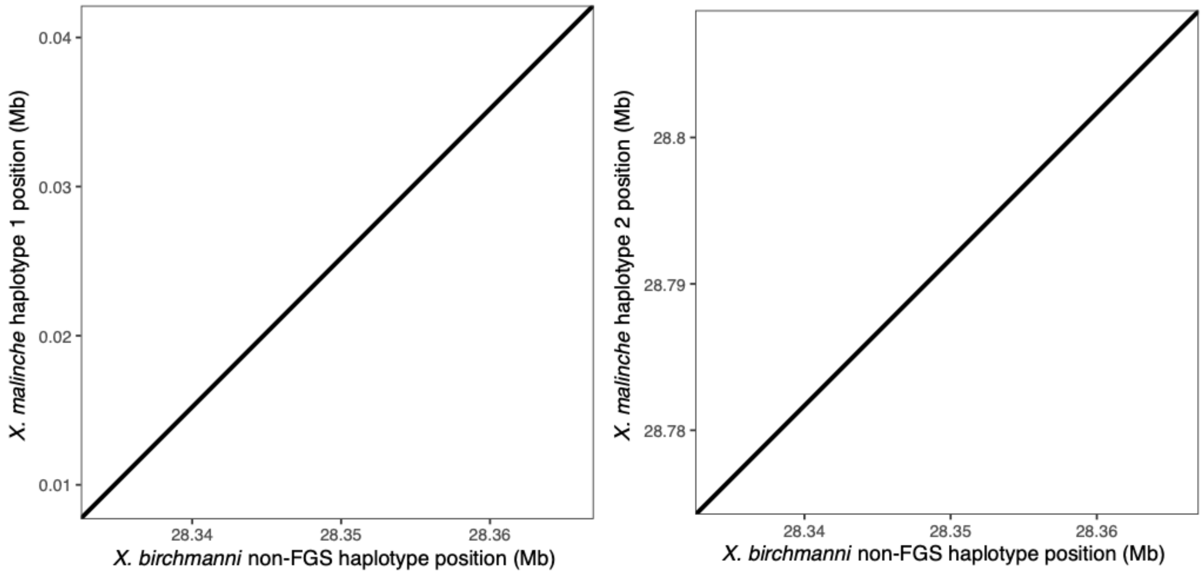


**Fig. S8.** MUMmer4 alignment of non-false gravid (non-FGS) spot *X. birchmanni* haplotype with both haplotypes from the *X. malinche* reference genome. *X. malinche* is fixed for the non-false gravid spot phenotype. Both haplotypes are completely co-linear with the non-false gravid spot *X. birchmanni* haplotype.

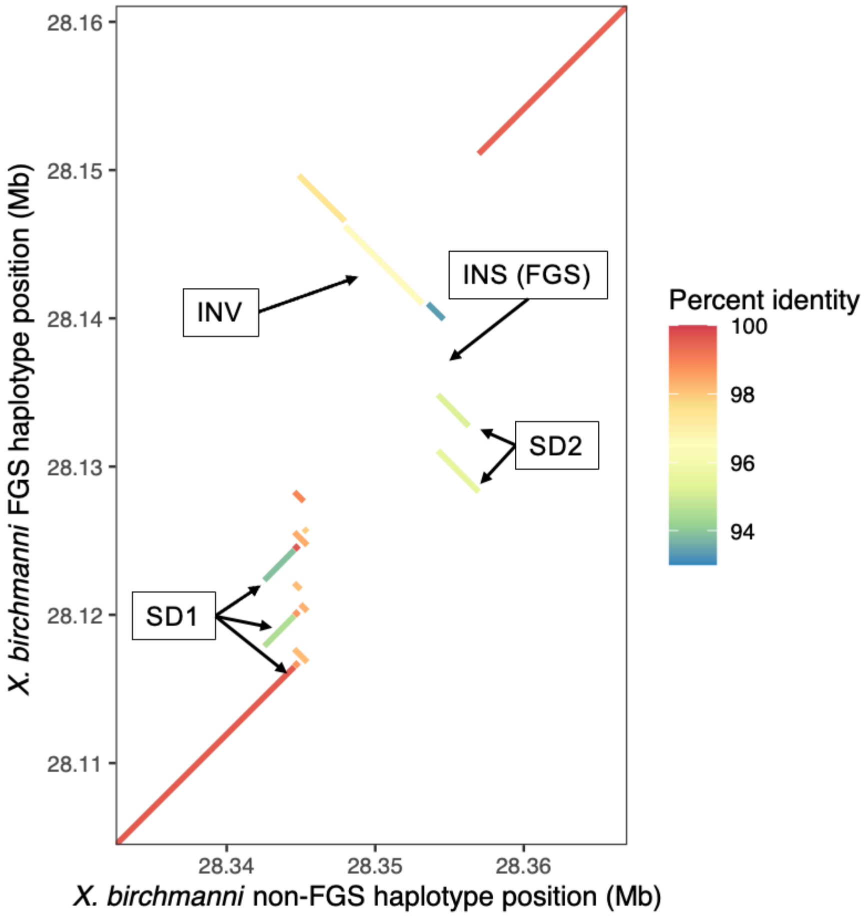


**Fig. S9.** MUMmer4 alignment of reference false gravid spot (FGS) and non-false gravid spot (non-FGS) haplotypes from *X. birchmanni*. Segments are colored by percent identity between the haplotypes. Individual components of the structural variant are labelled. SD1 - segmental duplication 1; SD2 - segmental duplication 2; INS (FGS) - insertion in false gravid spot haplotype (piggyback 4 element); INV - inverted region between false gravid spot and non-false gravid spot haplotype. INS (non-FGS) - insertion in non-false gravid haplotype (INDELs are visualized as a gap in one alignment or the other).

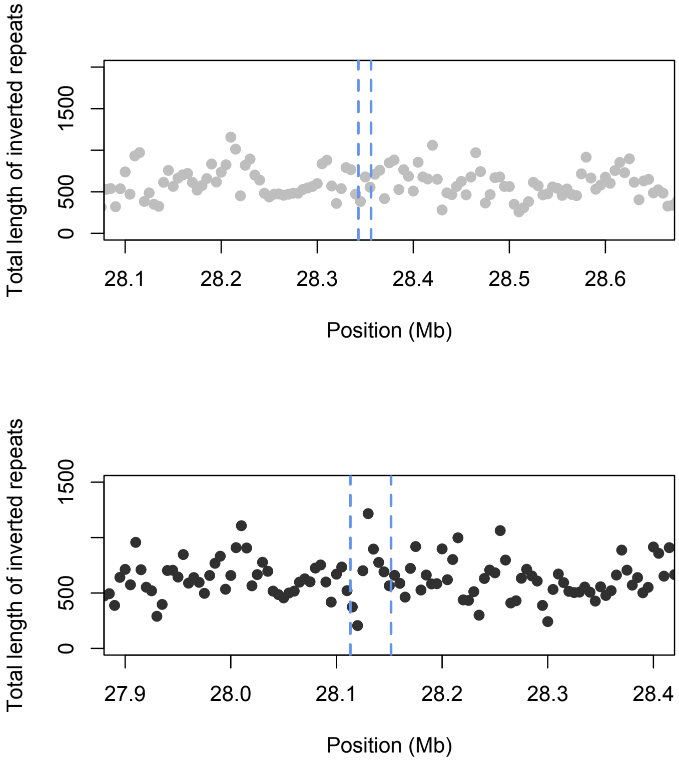


**Fig. S10.** Comparison of the length of inverted repeat sequences in sliding 5 kb windows identified by nBMST between the non-false gravid spot (top) and false gravid spot (bottom) haplotypes. Inverted repeats are enriched in the false gravid spot haplotype (bottom, black) compared to the non-false gravid spot reference sequence (top, grey). Significant regions in the GWAS analysis based on both reference haplotypes are noted by the blue dashed lines. Each dot represents the length in basepairs of the 5 kb region attributable to inverted repeats.

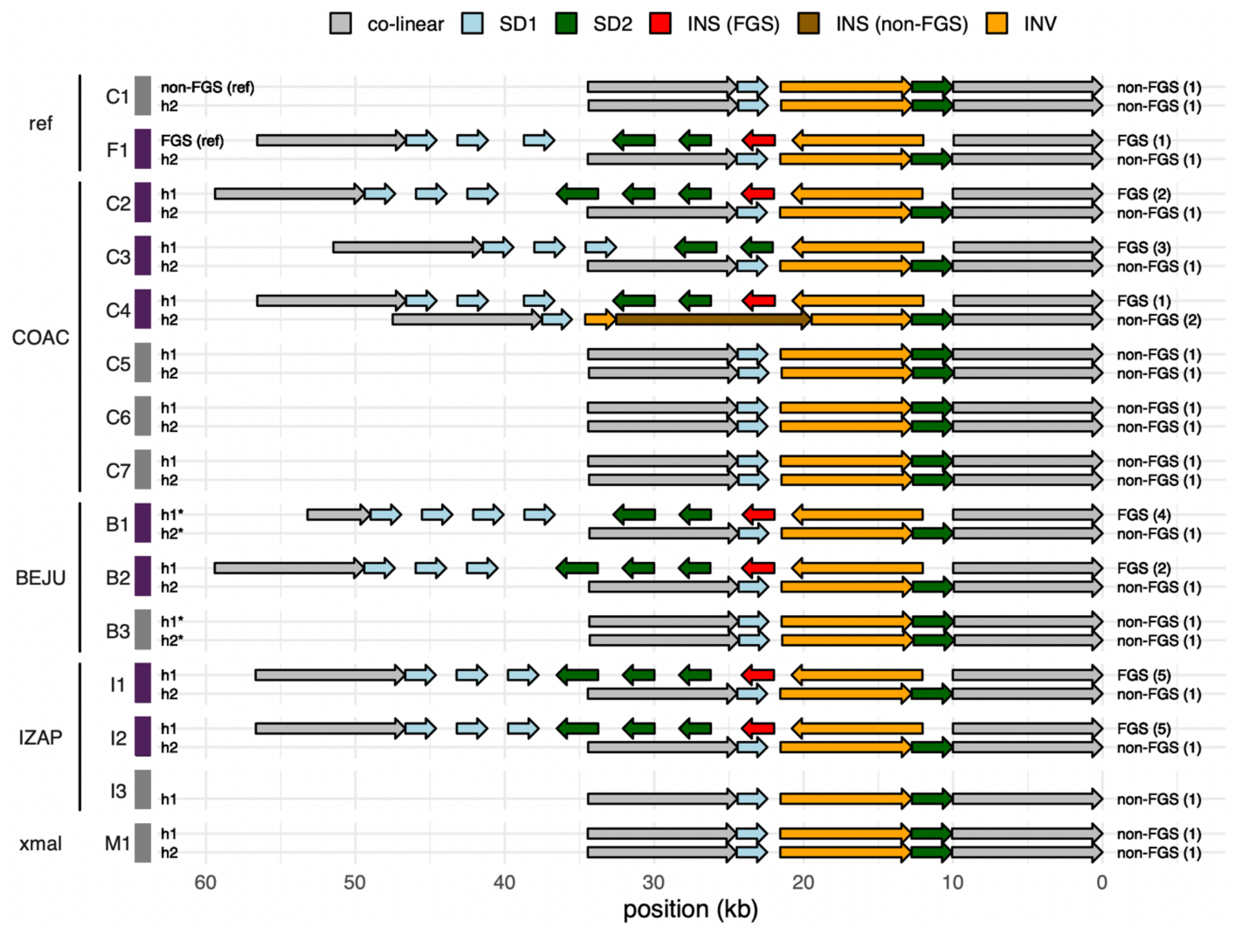


**Fig. S11.** Cartoons of structural variation in haplotypes generated for this study, based on MUMmer4 alignment coordinates for each individual. Haplotypes shown here are the same as Fig. 2b, with the addition of a pure *X. malinche* individual. Populations and species are denoted on left (reference individuals – ref; *X. birchmanni* Coacuilco – COAC; *X. birchmanni* Benito Juarez – BEJU; *X. birchmanni* Izapa – IZAP; *X. malinche* – xmal). Alpha numeric labels on left denote individuals, with the first letter typically corresponding to population, and the number denoting the sample (e.g. B1– Benito Juarez individual 1). Note that reference individual "ref F1" is an $F_1$ hybrid between *X. birchmanni* x *X. malinche*, so F1_h2 is a *X. malinche*-derived haplotype. Colors on left indicate false gravid spot (FGS) phenotype of the focal individual, with purple denoting false gravid spot and gray denoting non-false gravid spot phenotypes. All individuals with the false gravid spot phenotype have one rearranged haplotype compared to the reference genome, and one haplotype that is largely colinear. All individuals without the false gravid spot (including X. malinche) have 2 haplotypes colinear with the reference. SD1 - segmental duplication 1; SD2 - segmental duplication 2; INS (FGS) - insertion in false gravid spot haplotype (piggyback 4 element); INS (non-FGS) - insertion in non-false gravid haplotype. Labels on right correspond to haplotype class, with variant alleles numbered.

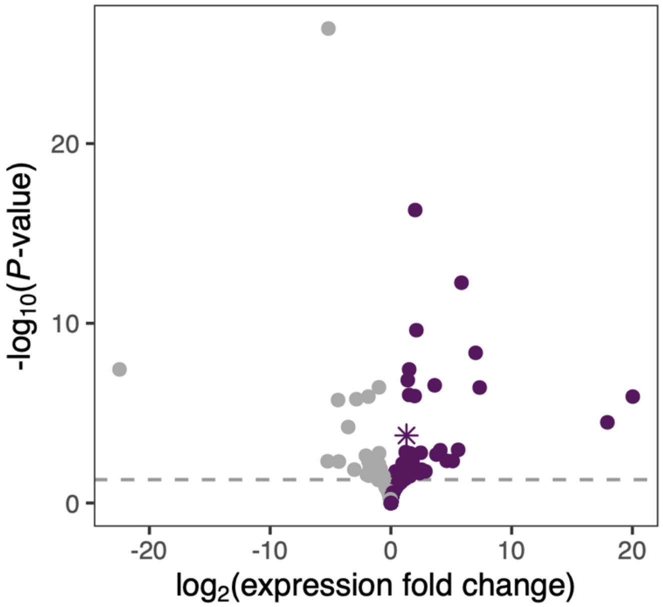


**Fig. S12.** Results of analysis of RNAseq data from the body wall musculature (flank) of false gravid and non-false gravid male *X. birchmanni*. The y-axis denotes $-\log_{10}$ of the false discovery rate adjusted p-value and the x-axis plots the $\log_2$ fold change in expression between groups. Points are colored based on if expression was increased in false gravid spot males (purple) or non-false gravid spot males (gray). *kitlga* is denoted with a star and was differentially expressed (LFC=1.30, $p<1.46e-7$). Dashed line corresponds to a false discovery rate adjusted p-value less than 0.05.

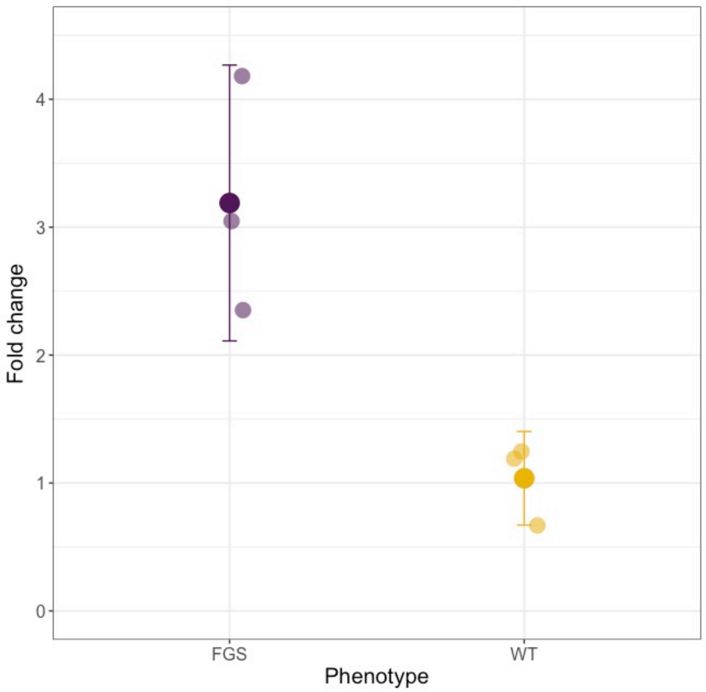


**Fig. S13.** Results of quantitative real-time PCR assay comparing expression of *kitlga* in PM + EAM tissue of false gravid males and non-false gravid spot makes. Fold change (delta delta ct) is compared to a housekeeping gene (*efa1*) and then standardized by mean non-FGS expression. Semi-transparent individuals points show biological replicates and larger points and whiskers show the mean ± 2 standard errors.

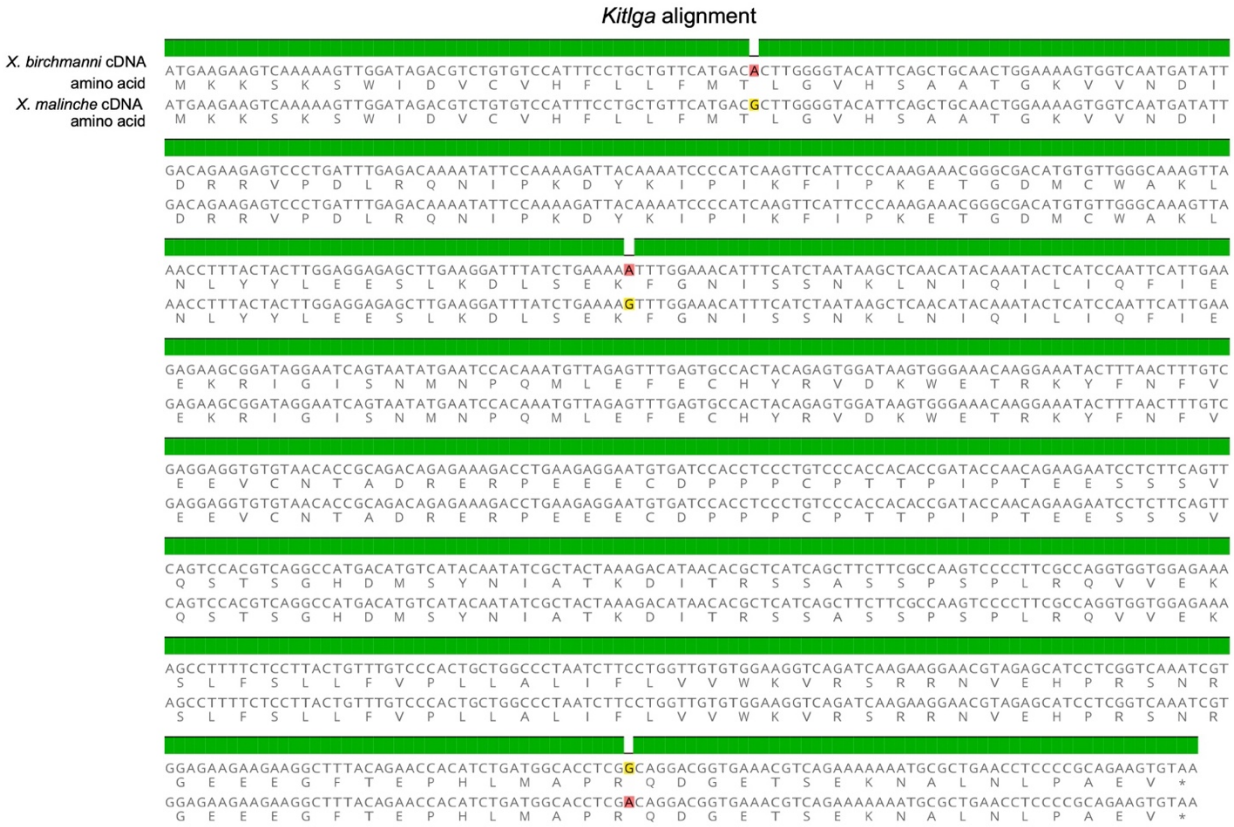

**Fig. S14.** Alignment of *kitlga* between *X. birchmanni* and *X. malinche* cDNA and amino acid sequences. Despite having no nonsynonymous substitutions, there are several fixed synonymous differences in the *kitlga* cDNA between *X. birchmanni* and *X. malinche.* These differences were used to design allele-specific expression primers.

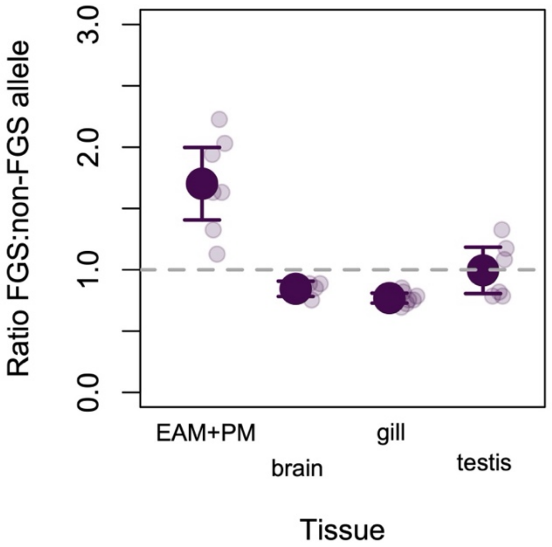


**Fig. S15.** Results from a second pyrosequencing primer targeting cDNA site 57 in *kitlga* confirm results presented in the main text that the false gravid spot allele is associated with higher tissue and allele specific expression in the EAM+PM tissue. The ratio of expression attributable to the false-gravid (FGS) versus non-false gravid (non-FGS) alleles is plotted on the y-axis. This suggests the expression differences in *kitlga* are under *cis*-regulatory control. Expression is normalized to the *X. malinche* allele. Large points and whiskers denote mean ± 2 standard errors, and small points represent individual results. Results shown in main text Figure 3 target cDNA site 759 in *kitlga*, which had slightly better performance in parental test samples.

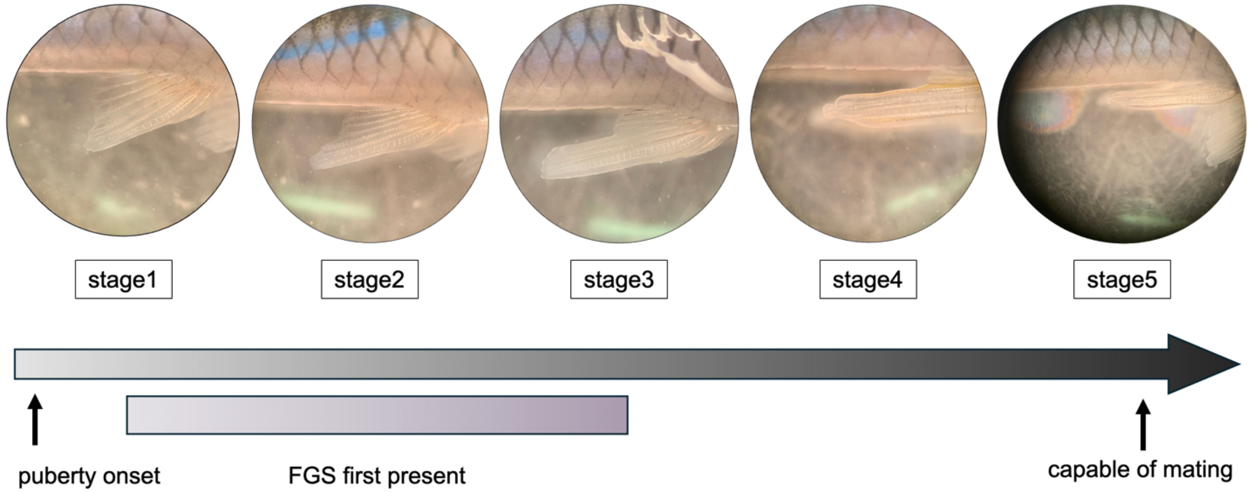


**Fig. S16.** Example images of gonopodial differentiation taken from the developmental time-tracking experiment. This experiment was used to determine when during gonopodial differentiation the false gravid spot phenotype developed. Males typically developed the false gravid spot phenotype between stages 1 and 3 of gonopodial development (indicated by the purple bar) but are not capable of mating until stage 5, when the hook and spikes form on the gonopodium and the cuticle recedes.

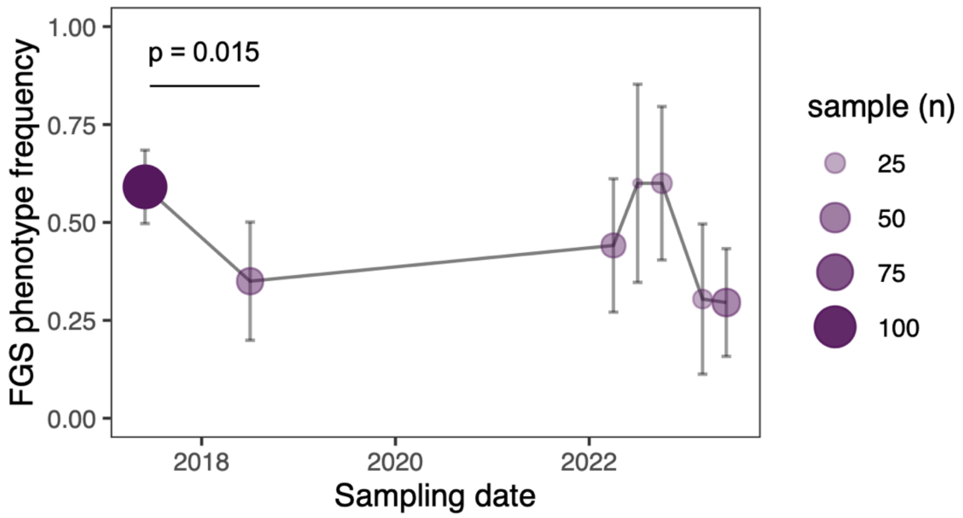


**Fig. S17.** Phenotypic frequencies of the false gravid spot (FGS) at Coacuilco fluctuate over time. Point size and opacity is proportional to the number of males collected at that time point. Error bars denote ± 2 binomial standard errors. Frequencies differed between 2017 and 2018 (two proportion Z-test, p=0.015).

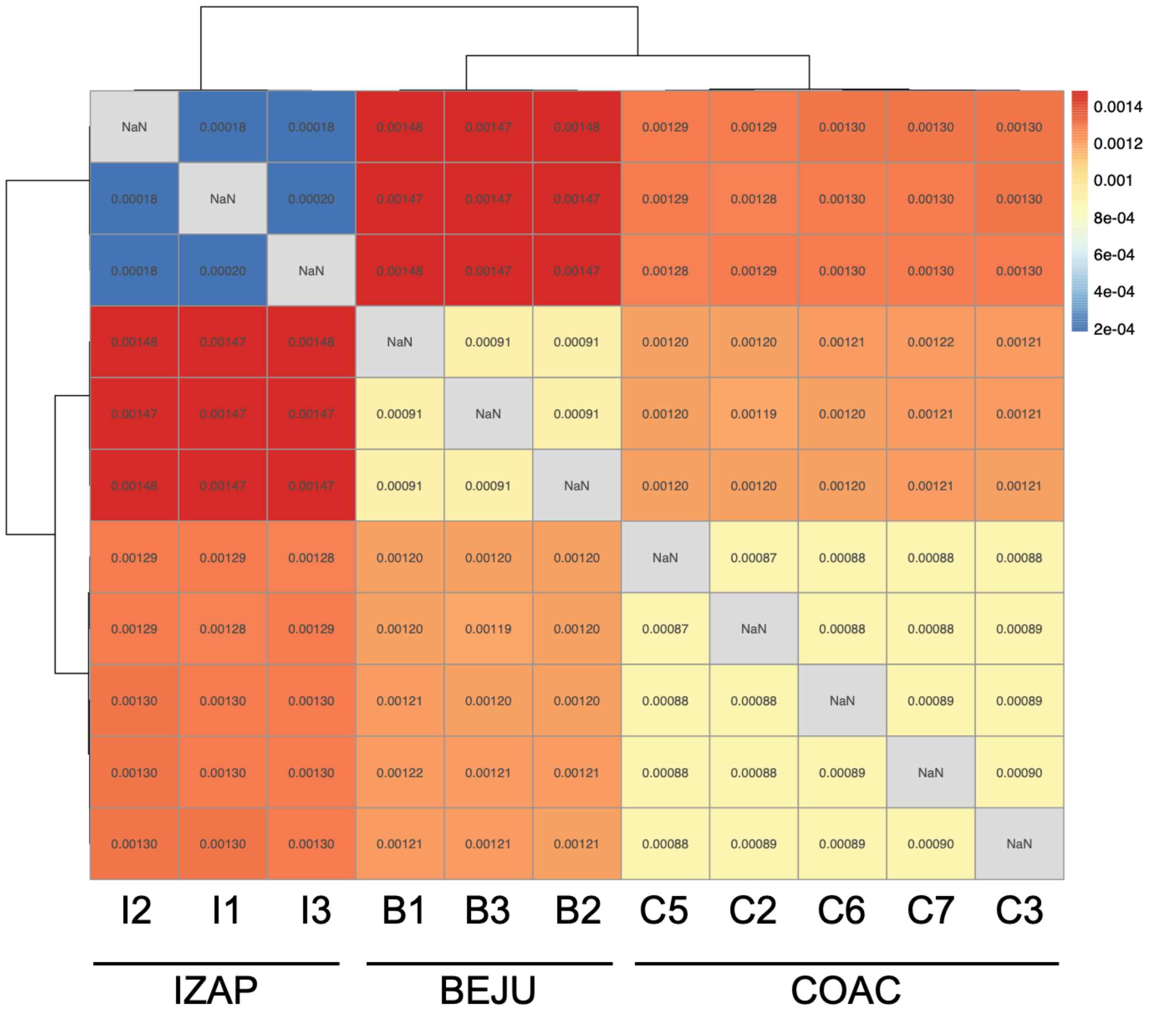


**Fig. S18.** Pairwise sequence divergence (dxy) between individuals across multiple *X. birchmanni* populations (IZAP – Izapa, BEJU – Benito Juarez, COAC – Coacuilco) sequenced with Oxford Nanopore. Individual labels are same as Fig. 2b. Dxy comparisons between individuals are plotted in pairwise matrix and organized using the clustering algorithm implemented in pheatmap. Dxy values between samples are labelled and colored, with warm colors denoting higher dxy values and blue denoting lower dxy values. This analysis suggests some signals of genetic divergence between distinct *X. birchanni* populations and also highlights variation in rates of polymorphism within populations.

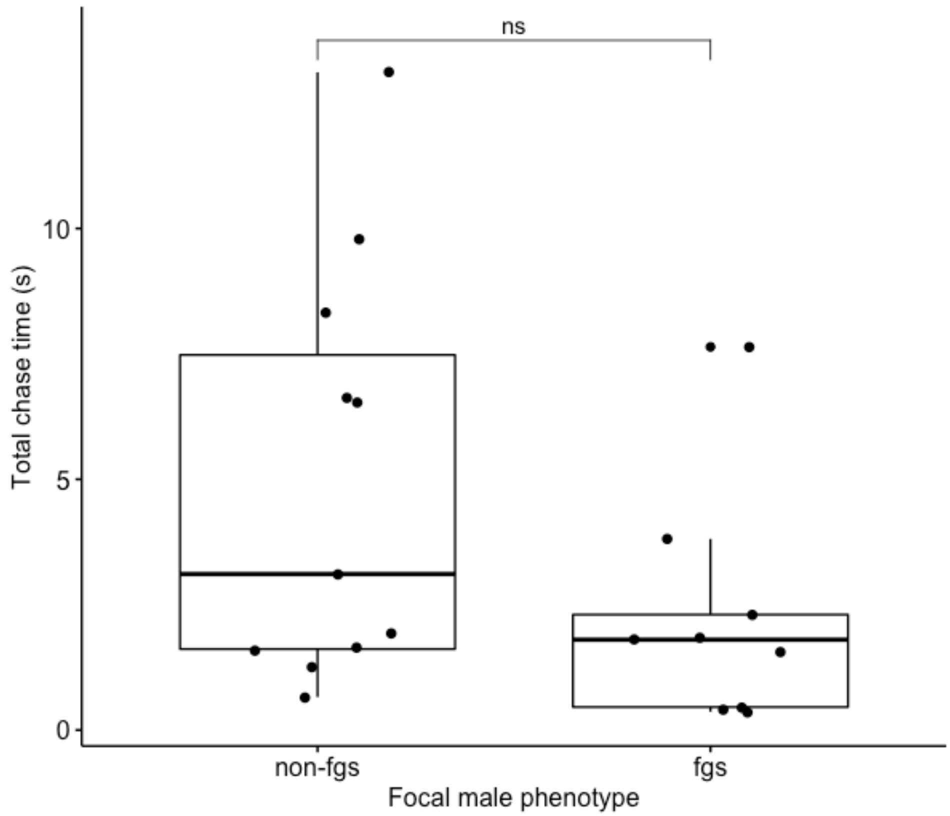


**Fig. S19.** Focal males did not differ in aggression based on their false gravid (FGS) phenotype as measured by total time spent chasing all targets (two-sample Welch's t-test; $t = 1.84$, $p = 0.085$). All other metrics of aggression yielded similar results. Points represent means across all trials for each focal male.

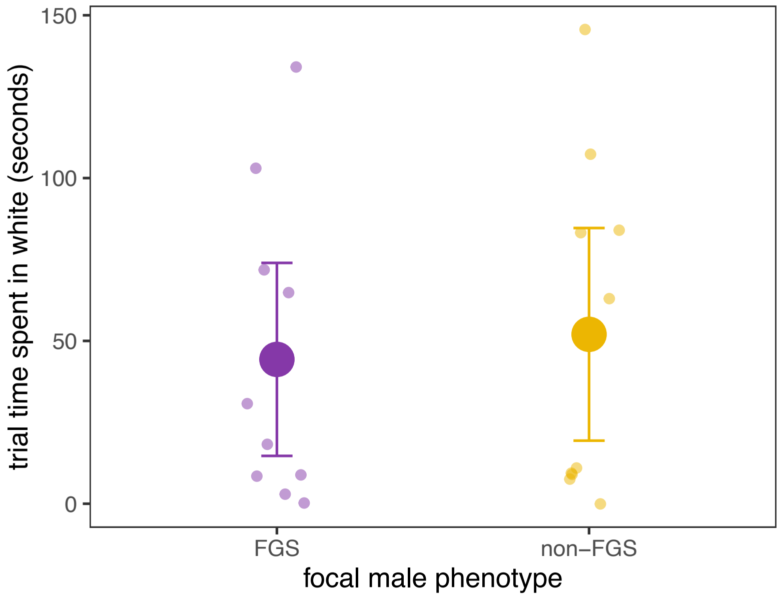


**Fig. S20.** Results of scototaxis trials to assay boldness in males with and without the false gravid spot. We found no difference in time spent on a white background between males with and without the false gravid spot trait (p=0.77), suggesting no clear difference in boldness between phenotypes in this setting. FGS – false gravid spot, non-FGS – non-false gravid spot. Small points indicate results of individual trials, large points and error bars indicate the mean ± 2 standard errors.

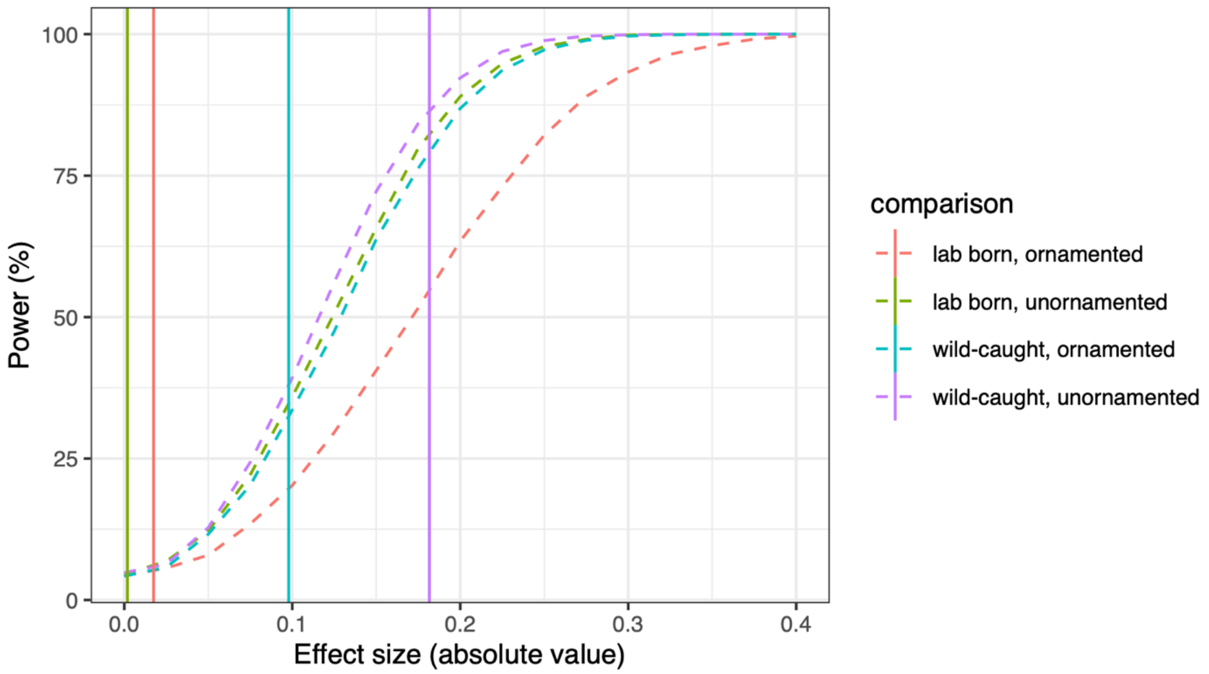


**Fig. S21.** Expected power to detect a true effect of varying strength in behavioral trials. Results of simulations for each female group and animation stimulus under scenarios with known effect sizes for female preference. Empirically inferred effect sizes (average difference in the proportion of the trial spent associated with each stimulus) from experimental data are shown with vertical dashed lines. Results indicate that we expect to have good power to detect large effect sizes, such as those detected in wild-caught females shown animations of unornamented males with and without the false gravid spot.

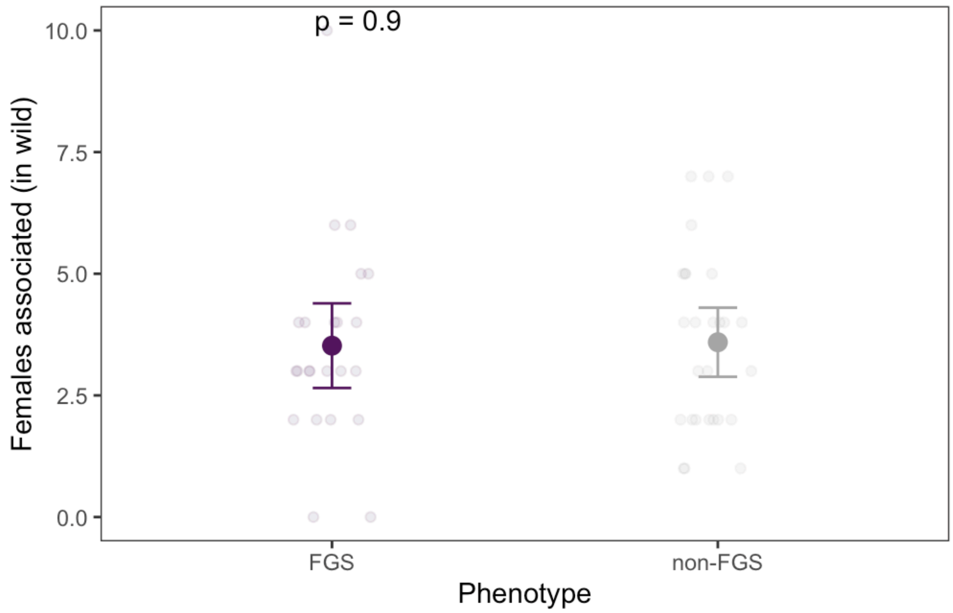


**Fig. S22.** In observations of males in the Coacuilco population, males with false gravid spot (FGS) and without false gravid spot (non-FGS) did not differ in the number of females found nearby (t-test; p=0.9). Dark points denote means. Error bars indicate ± 2 standard errors. Light points indicate individual observations.

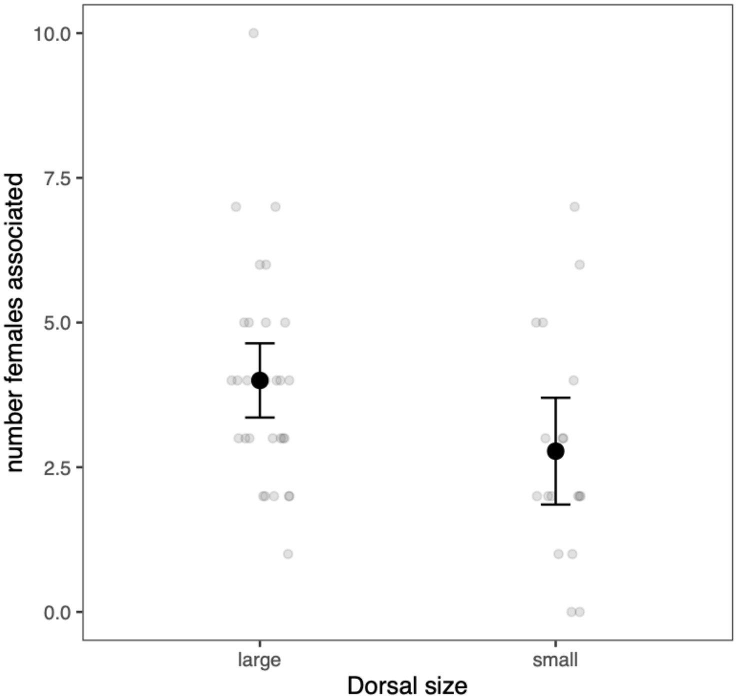


**Fig. S23**. Variation in the mean number of females found in association with the focal males was best explained by the size of a male's dorsal fin. GLM: Likelihood ratio $\chi^2_1 = 4.9$, P = 0.027. Dark points denote means. Error bars indicate ± 2 standard errors. Light points indicate individual observations.

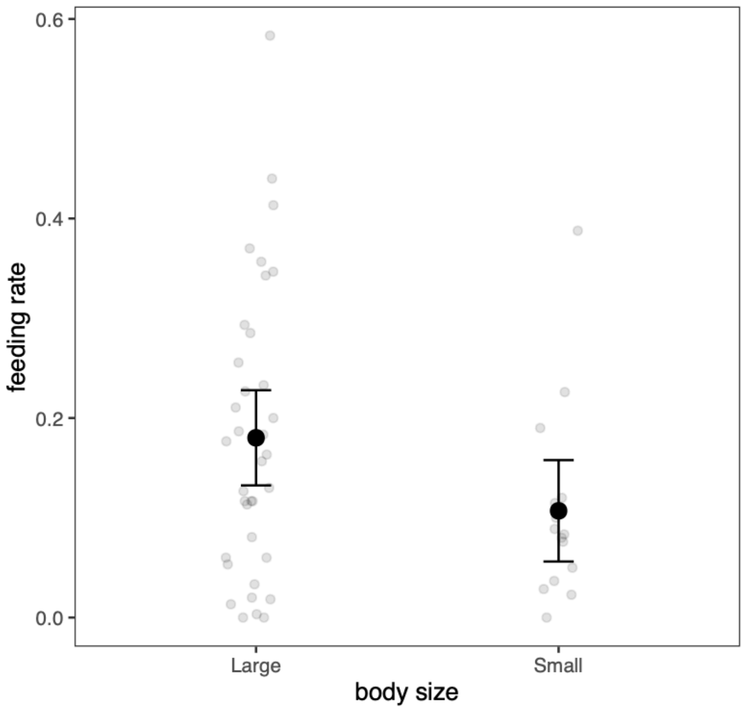


**Fig. S24.** Variation in male feeding rate (nips at substrate per minute) is best explained by male size in observations in a natural population. Large body size was defined as males longer than a 4.4 cm and small body size was defined as males under 4.4 cm. This value was chosen because it corresponds to the mean male body size at Coacuilco (GLM: Likelihood ratio $\chi^2_1 = 4.4$, $P = 0.036$). Dark points denote means. Error bars indicate ± 2 standard errors. Light points indicate individual observations.

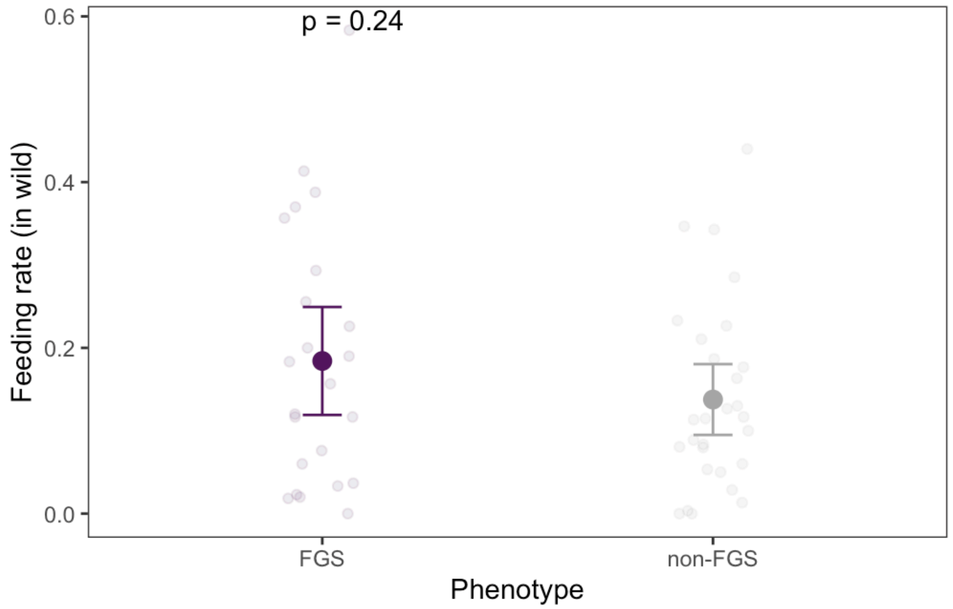


**Fig. S25.** In observations of males in the Coacuilco population, males with and without false gravid spot did not differ in their feeding rates (GLM: Likelihood ratio $\chi^2_1 = 2.3$, $P = 0.132$). Dark points denote means. Error bars indicate ± 2 standard errors. Light points indicate individual observations. FGS – false gravid spot males, non-FGS – non- false gravid spot males.

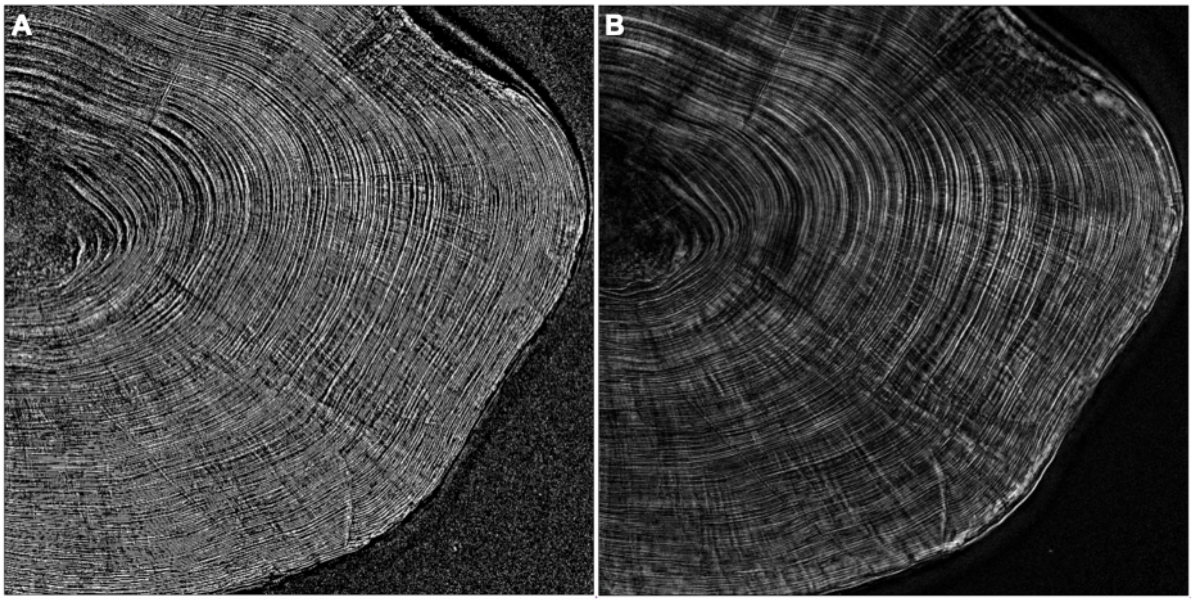


**Fig. S26.** Example images from otolith image and data processing. We post-processed the image stacks in two ways, shown here. **A)** We compiled a max projection of each image stack and sharpened once and enhanced contrast by 0.35%. **B**) Second, we selected a single image from each stack and sharpened it once. Otolith rings were counted by 4 observers.

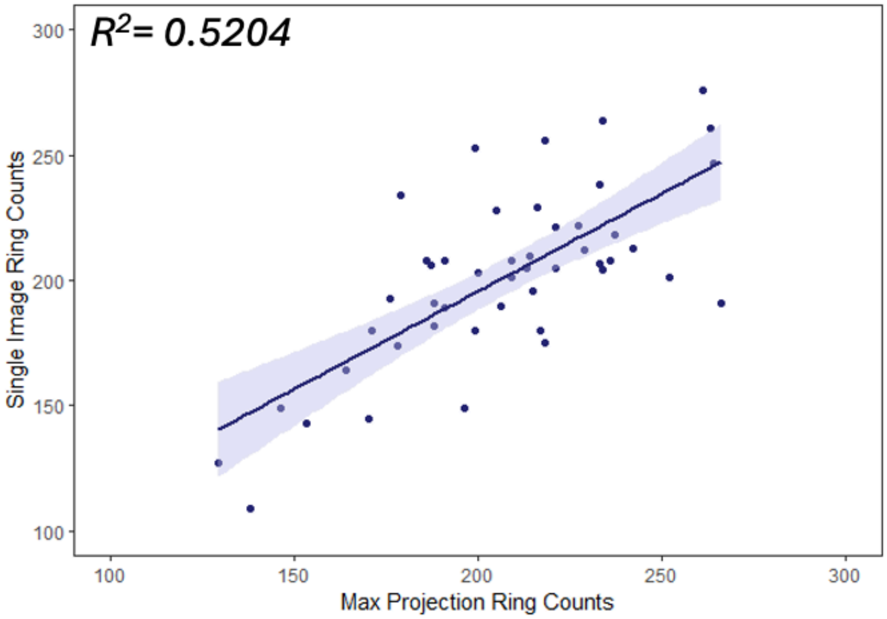


**Fig. S27.** Manual counts on max projection images of otoliths compared to single images yielded similar ring counts ($p<0.001$). However, the relationship was weaker than expected given that these results are technical replicates derived from counts from the same otolith. For subsequent analyses, the mean of the two counting methods was used.

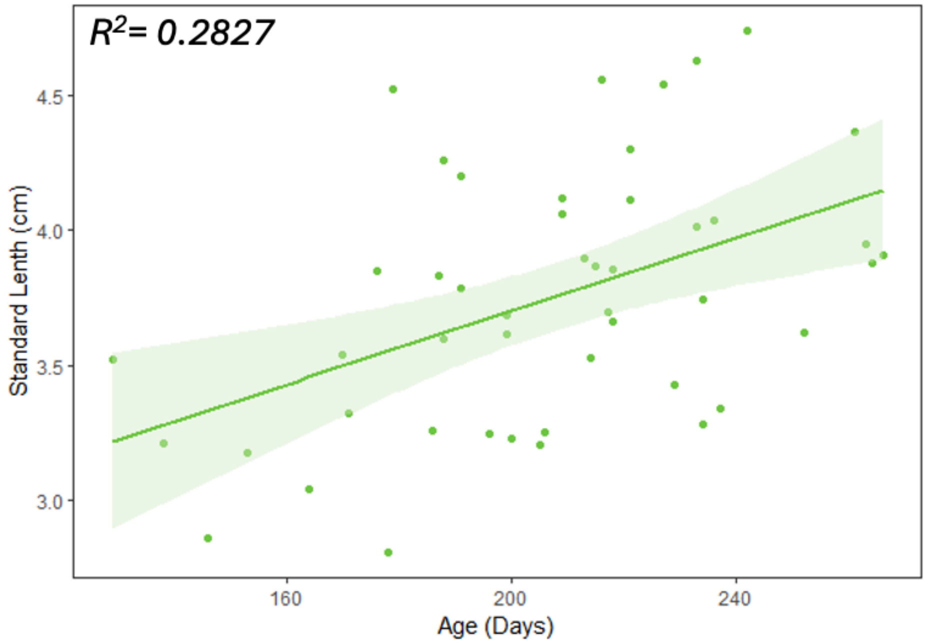


**Fig. S28.** Male standard length is strongly associated with otolith ring count in our dataset ($p<0.0007$). This result is expected as males increase in size up until sexual maturity and also lay down separable daily otoliths up until sexual maturity.

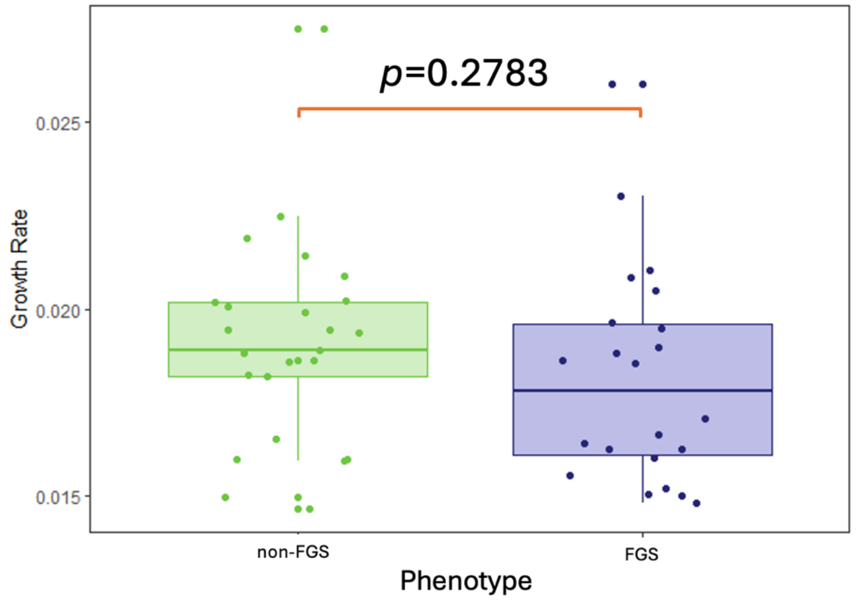


**Fig. S29.** Juvenile male *X. birchmanni* with the false gravid spot (FGS) and fish without false gravid spot (non-FGS) do not have significantly different growth rates (p<0.28) in samples from natural populations. Growth rates were calculated by dividing standard length by otolith ring count (expected to be equivalent to age in days).

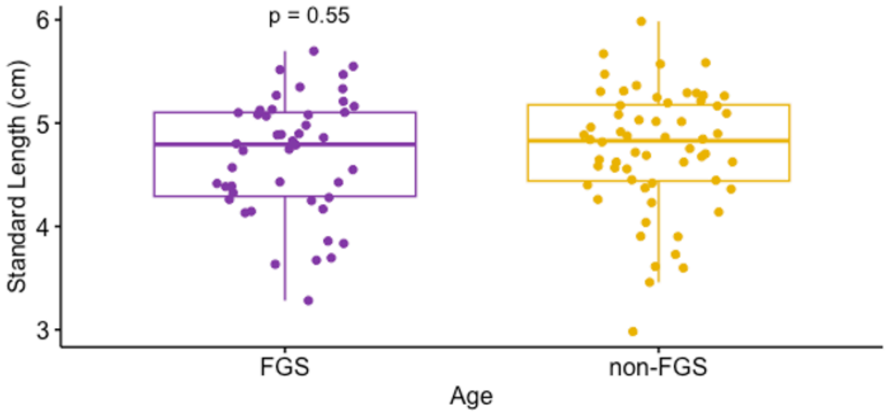


**Fig. S30.** Standard length in adult male *X. birchmanni* from Coacuilco collected between 2017 and 2018. Males with and without false gravid spot did significantly differ in standard length. FGS – males with false gravid spot; non-FGS – males without false gravid spot.
